# Sigh generation in preBötzinger Complex

**DOI:** 10.1101/2024.06.05.597565

**Authors:** Yan Cui, Evgeny Bondarenko, Carolina Thörn Perez, Delia N. Chiu, Jack L. Feldman

## Abstract

We explored neural mechanisms underlying sighing. Photostimulation of parafacial (pF) neuromedin B (NMB) or gastrin releasing peptide (GRP), or preBötzinger Complex (preBötC) NMBR or GRPR neurons elicited ectopic sighs with latency inversely related to time from preceding endogenous sigh. Of particular note, ectopic sighs could be produced without involvement of these peptides or their receptors in preBötC. Moreover, chemogenetic or optogenetic activation of preBötC SST neurons induced sighing, even in the presence of NMBR and/or GRPR antagonists. We propose that an increase in the excitability of preBötC NMBR or GRPR neurons not requiring activation of their peptide receptors activates partially overlapping pathways to generate sighs, and that preBötC SST neurons are a downstream element in the sigh generation circuit that converts normal breaths into sighs.

## Introduction

Endogenous sighs generated periodically, typically every few minutes in rodents and humans, reinflate collapsed alveoli to maintain proper gas exchange in the mammalian lung^1–3^. Two largely parallel medullary pathways from the parafacial region (pF) to preBötzinger Complex (preBötC) appear critical for generation of these physiological sighs: pF neurons expressing neuromedin B (NMB) or gastrin releasing peptide (GRP) project to preBötC neurons expressing their cognate receptors, NMBR and GRPR^4^. These pathways also appear to mediate confinement-induced claustrophobic sighing^5^, and by extension, other forms of sighing associated with emotional states such as during sadness, anxiety, depression, relief, or happiness^6,7^.

In rodents, the preBötC receives inputs from NMB and GRP pF neurons, and genetic deletion of NMBRs or GRPRs or local antagonism of NMBRs or GRPRs in preBötC significantly reduces endogenous sighing, while local injection into preBötC of bombesin, NMB or GRP significantly increases spontaneous sighing^4^. In order to determine whether NMB and GRP are obligatory for sighs, and to better understand the mechanisms within the preBötC underlying sigh generation, we investigated whether activation of pF NMB or GRP neurons can generate sufficient input to preBötC to induce sighs and whether activation of preBötC GRPRs or NMBRs is necessary for generation of sighs.

Using transgenic mice that express Cre- or Flp-recombinase in *Nmb*-, *Grp*-, *Nmbr*- or *Grpr*- expressing neurons, we investigated the effects on sighing and other aspects of breathing pattern *in vivo*: i) of optogenetic activation of ChR2-transfected pF GRP or NMB neurons; ii) of optogenetic or chemogenetic activation of ChR2-transfected preBötC GRPR or NMBR neurons, including in the presence of their antagonists, and; iii) determined whether activating preBötC SST neurons, presumptive preBötC output neurons, can generate sighs. Moreover, by using electrophysiology and two-photon calcium imaging, we investigated the activity profiles of *Grpr*- or *Nmbr*-expressing preBötC neurons *in vitro*.

We conclude that: i) activation of either pF NMB or GRP neurons generates sufficient input to preBötC to generate ectopic sighs; ii) sighs can be generated by activation of preBötC GRPR and NMBR neurons even after blockade of NMBR and GRPR receptors; iii) preBötC GRPR or NMBR neurons act via partially overlapping pathways to produce sighs via preBötC SST neurons; iv) preBötC NMBR neurons are not rhythmogenic, and; v) activation of preBötC SST neurons can generate sighs.

## Results

A sigh is an inspiratory effort that results in significantly increased tidal volume (V_T_), typically two to five times larger compared to normal breaths with a range of airflow profiles. Here, we define sighs as inspiratory efforts that result in transient significantly (>2-fold) increased V_T_, which occur periodically at intervals longer than eupneic, i.e., normal, breaths. For all analyses, we designated sighs by their larger V_T_ irrespective of their shape. Thus, monophasic ‘augmented breaths’, biphasic ‘partial doublets’ or ‘full doublets’ were all considered sighs (Extended Data Fig. 1). The mechanisms underlying formation of these different sigh subtypes are outside the scope of this publication.

### Activation of *Nmb*- or *Grp*-expressing pF neurons induces sighs

To test whether selective activation of pF NMB or GRP neurons, i.e., bombesin peptide-expressing pF neurons) generates sighs, we established Grp-ChR2 or Nmb-ChR2 mouse lines (see *Methods*) by crossing floxed-ChR2-tdTomato mice with Grp^Cre^ and Nmb^Cre^ mice (Fig. 1a, c). We then measured the effects of targeted pF photostimulation (long pulse photostimulation, LPP, single bilateral 4-10 s 5mW pulse or short pulse photostimulation, SPP, single bilateral 100-500 ms pulse; see *Methods*)^8^ in anesthetized Grp-ChR2 or Nmb-ChR2 mice.

**Fig. 1.**
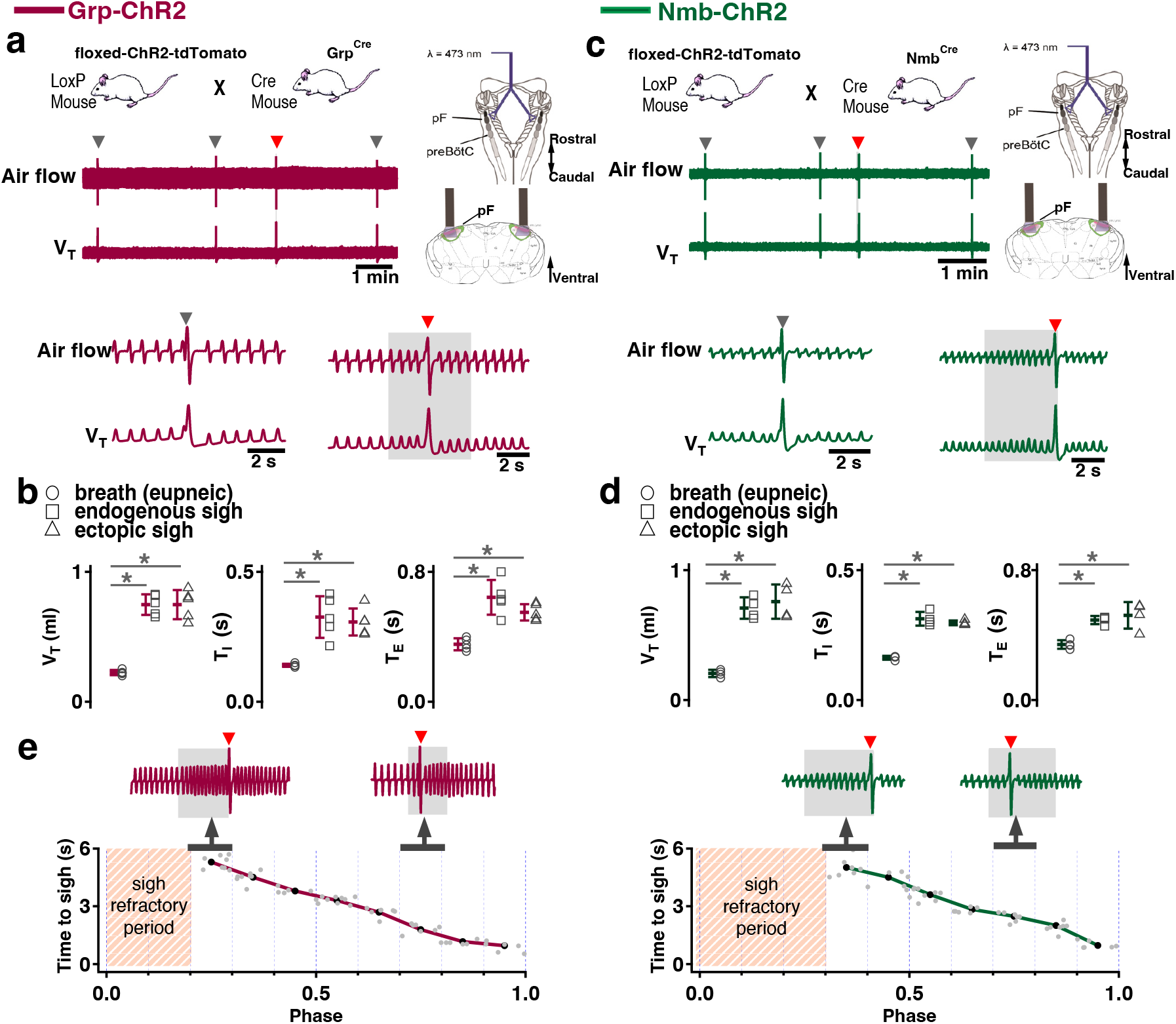
**Photostimulation of NMB and GRP pF neurons evoked sighing**. **a**, Top: Breeding scheme to generate Grp-ChR2 mice. Schematic (right) depicting bilateral placement of optical cannula targeting pF. Middle and Bottom: Raw (middle) and expanded (bottom) traces show bilateral pF LPP (right gray box) in Grp-ChR2 mice elicits an ectopic sigh (red arrowhead) that appears similar to endogenous sighs (gray arrowheads). **b**, V_T_ (t_3_ = 0.920, p = 0.393), *T_I_* (t_3_ = 1.811, p = 0.120), and *T_E_* (t_3_ = 0.277, p = 0.791) of ectopic sighs from pF Grp-ChR2 photostimulation were no different from endogenous sighs (n = 4 mice), but increased compared to eupneic breaths (V_T_: t_4_ = 10.163, p = 5 x 10^-5^ *T_I_*: t_3_ = 6.224, p = 7 x 10^-4^, *T_E_*: t_4_ = 7.152, p = 4 x 10^-4^), indicative of augmented breaths with postsigh apneas. Statistical significance was determined with a One Way RM ANOVA followed by All Pairwise Multiple Comparison Procedures (Holm-Sidak method), V_T_: F_2,9_ = 63.194, p < 0.001; *T_I_*: F_2,9_ = 35.529, p < 0.001; *T_E_*: F_2,9_ = 32.828, p < 0.001. **c**, Top: Breeding scheme to generate Nmb-ChR2 mice. Schematic (right) depicting bilateral placement of optical cannula targeting pF. Middle and Bottom: Raw (middle) and expanded (bottom) traces show bilateral pF LPP (right gray box) in Nmb-ChR2 mice elicits ectopic sighs (red arrowhead) that appears similar to endogenous sighs (gray arrowheads). **d**, V_T_ (t_3_ = 2.436, p = 0.0508), *T_I_* (t_3_ = 0.603, p = 0.569), and *T_E_* (t_3_ = 0.308, p = 0.768) of ectopic sighs from pF Nmb-ChR2 photostimulation were no different from endogenous sighs (n = 4 mice), but increased compared to eupneic breaths (V_T_: t_3_ = 37.257, p = 3 x 10^-8^, *T_I_*: t_3_ = 6.697, p = 5 x 10^-4^, *T_E_*: t_3_ = 7.074, p = 4 x 10^-4^), indicative of augmented breaths with postsigh apneas. Statistical significance was determined with a One Way RM ANOVA followed by All Pairwise Multiple Comparison Procedures (Holm-Sidak method), V_T_: F_2,9_ = 868.829, p < 0.001; *T_I_*: F_2,9_ = 27.450, p < 0.001; *T_E_*: F_2,9_ = 34.883, p < 0.001. **e**, Sigh latency from laser onset was negatively correlated with sigh phase in Grp-ChR2 (r = -0.995, p = 3 x 10^-7^) and Nmb-ChR2 (r = -0.993, p = 8 x 10^-6^) mice (Pearson Product Moment Correlation). Top: Representative traces showing that latency between laser onset (gray box indicates LPP) and ectopic sighs (red arrowheads) was longer when LPP was applied just after the refractory period (pink diagonal stripes). Gray dots represent latency to ectopic sigh from stimulation onset in each phase in Grp-ChR2 (n = 5 mice) and Nmb-ChR2 (n = 4 mice) mice, solid lines join the average latency in each phase (black circle). No sighs were generated by LPP in sigh phase 0.0–0.2 (34 ± 5 s from previous sigh, measured from 30 stimuli in 5 mice: basal sigh rate 21 ± 3/hour, range 17–25; intersigh interval 172 ± 25 s) in Grp-ChR2 and in phase 0.0–0.3 (58 ± 20 s from previous sigh, measured from 48 stimulus in 4 mice: basal sigh rate 20 ± 7/hour, range 13-27; intersigh interval 193 ± 65 s) in Nmb-ChR2 mice. Data are shown as mean ± SE. Asterisks indicate post-hoc multiple comparison test results or paired t-test results: *, significance with p < 0.05.

In brainstems of adult Grp-ChR2 mice at the rostrocaudal level of the facial nucleus (7N), tdTomato fluorescence is expressed mainly in the dorsomedial part of retrotrapezoid nucleus/parafacial respiratory group (RTN/pFRG; Extended data Fig. 2a), consistent with previous data in newborn mice^4^. In brainstems of adult Nmb-ChR2 mice, Nmb-tdTomato is mainly expressed ventral and lateral to 7N, with a few of these neurons in the 7N, consistent with previous data in mice^9^ (Extended data Fig. 2b).

**Fig. 2.**
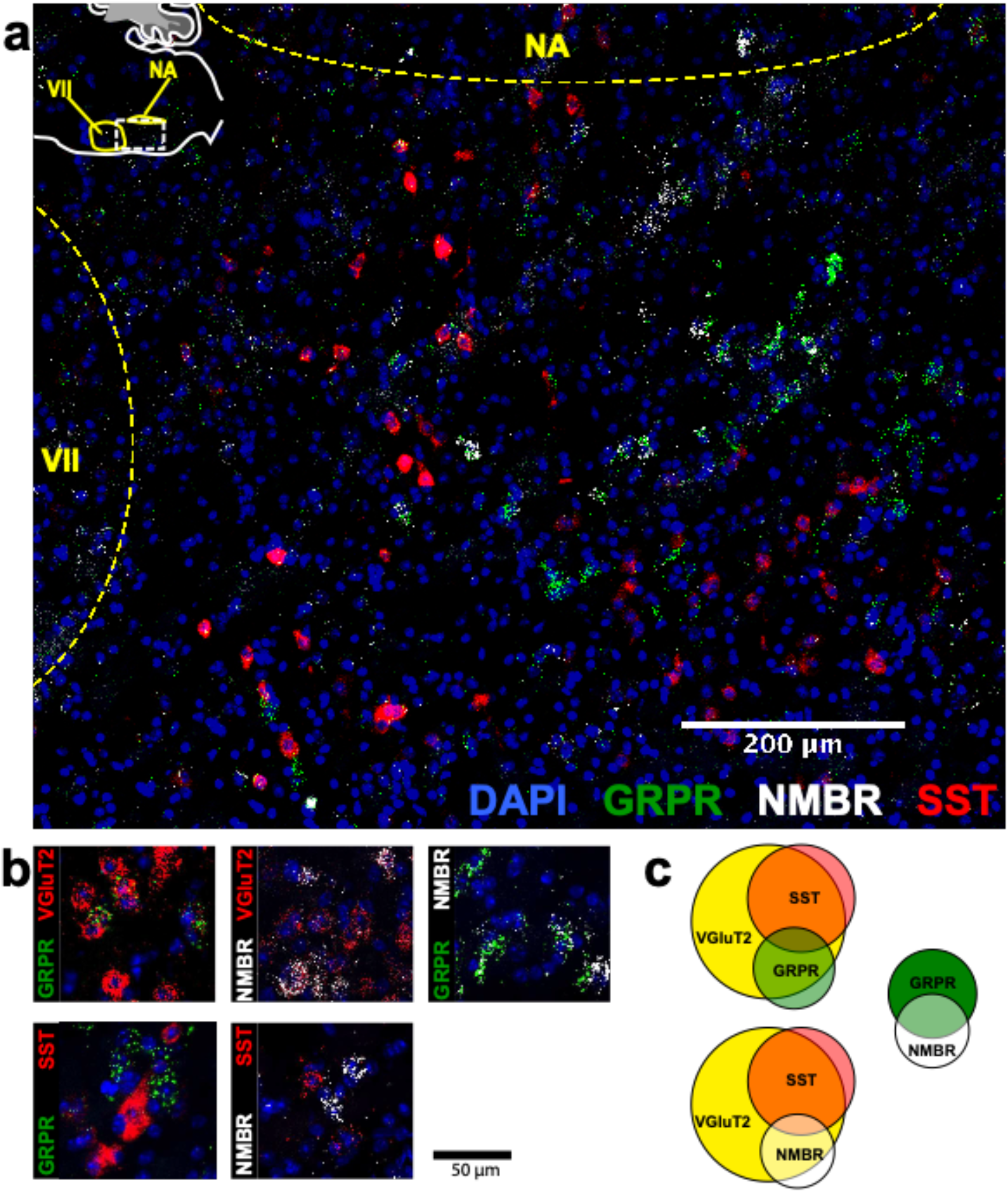
NMBR and GRPR neurons in preBötC are mainly glutamatergic, but mostly not somatostatinergic. **a**, Sagittal section of medulla with NMBR (white), GRPR (green) and SST (red) RNAscope signals as well as DAPI (blue) at the level of preBötC; inset top left: location of image (line box) in mouse medulla. **b**, Examples of colocalization between relevant markers in high magnification confocal images (100 x 100 μm) of tissues processed with RNAscope. **c**, Venn diagrams representing relative number of preBötC neurons expressing relevant markers and their overlap, scaled according to total Vglut2 count (n = 3 sections for each colocalization pair). VII: facial nucleus. NA: nucleus ambiguous. There were 22.6 ± 2.3 GRPR^+^ neurons and 14.6 ± 1.4 NMBR^+^ neurons in each transverse section; 25/45 NMBR^+^ neurons coexpressed GRPR; 25/77 GRPR^+^ neurons coexpressed NMBR. Majority of both NMBR^+^ and GRPR^+^ co-expressed VgluT2 (47/55 and 63/76, respectively), but not SST (6/46 and 5/54, respectively).

In Grp-ChR2 mice, LPP in pF elicited a large breath equivalent to an endogenous sigh in both amplitude and tidal volume (V_T_) when initiated during phase 0.3-1.0 of the sigh cycle (see *Methods*); we refer to such events as ectopic sighs (Fig. 1a, bottom). Ectopic sighs in Grp-ChR2 mice were of partial doublet or augmented breath shape (See Extended Data Fig.1) and were indistinguishable from endogenous sighs in both V_T_ and inspiratory (*T_I_*) and expiratory (*T_E_*) duration and exhibited a postsigh apnea (*T_E_* immediately following ectopic sighs increased to 216 ± 35% of *T_E_* of eupneic breaths, Fig.1b). During sigh phase 0.3-0.9, the interval from the previous endogenous sigh to the LPP-induced ectopic sigh, i.e., the perturbed sigh cycle, was shorter than the prior control sigh cycle. The interval between an ectopic sigh and the next (endogenous) sigh was, on average, indistinguishable from the interval between endogenous sighs, indicating that ectopic sighs reset the sigh cycle (Fig. 1a). No ectopic sighs could be generated by bilateral pF SPP in Grp-ChR2 mice during inspiration or expiration (phase: 0.0- 1.0).

In Nmb-ChR2 mice, the effects of pF SPP and LPP were similar to those in Grp-ChR2 mice, as SPP (phase: 0.0-1.0) had no effect on sighing while LPP induced ectopic sighs, which appeared as partial doublet or augmented breaths, and reset the sigh cycle during 0.4-0.9 sigh phase (Fig 1c). Ectopic sighs were also indistinguishable from spontaneous sighs in V_T_, *T_E_* and *T_I_* and they exhibited a postsigh apnea (*T_E_* after evoked sighs increased to 151 ± 22% of *T_E_* of eupneic breaths, Fig. 1d).

Notably, in either Grp-ChR2 or Nmb-ChR2 mice, pF LPP did not elicit ectopic sighs when applied very shortly after an endogenous sigh (phase range: 0.0–0.2 and 0.0–0.3 of endogenous sigh cycle, respectively; Fig. 1e), indicating a postsigh refractory period for sigh induction. LPP in these cases had no subliminal effect on the sigh cycle, as the expected time to the next endogenous sigh was unchanged.

When pF LPP was initiated immediately following the end of the postsigh refractory period, the perturbed sigh cycle was significantly shorter than the control sigh cycle. For pF LPP later in the sigh cycle, i.e., closer to when the next endogenous sigh was expected, ectopic sighs occurred at even shorter latencies (Fig. 1e). Thus, after the refractory period during which ectopic sighs could not be elicited, this inverse relationship between photostimulation phase and sigh latency indicated that acute activation of the peptidergic microcircuit was more effective as the sigh phase advanced.

### NMBR and GRPR are expressed primarily in excitatory neurons in preBötC

To understand how the activity of peptide-expressing pF neurons results in the generation of sighs, we next examined expression pattern of *Nmbr*- and *Grpr*- expressing cells in the preBötC using fluorescence *in situ* hybridization (FISH; Fig. 2). The vast majority of NMBR and GRPR cells (>97%) did not colocalize with the specific astrocytic marker Aldh1l1, indicating that these cells are neurons (Extended Data Fig. 3). GRPR neurons were present caudal to pF, in a stream continuing dorsocaudally from pF to preBötC. In contrast, with the exception of sparse NMBR/ChAT neurons in nucleus ambiguus (Amb. ∼4 neurons in each section) that were excluded from subsequent analysis, NMBR neurons were mainly found in preBötC. As previously reported^4^, some neurons coexpressed *Nmbr* and *Grpr* (Fig. 2b). In our sample, 57% of NMBR neurons coexpressed *Grpr*, which constituted 32% of the GRPR population (Fig. 2b, c).

Next, we assessed colocalization with the vesicular glutamate transporter VGlut2 to determine whether NMBR and GRPR neurons are glutamatergic. Approximately 85% of NMBR and GRPR neurons co-expressed VGlut2 (Fig. 2b, c). Finally, we asked whether *Nmbr* and *Grpr* were expressed on preBötC SST neurons. We found scant colocalization of SST and NMBR or GRPR (∼10%), indicating that the substantial majority of these peptide receptors are not on SST neurons (Fig. 2b, c).

### Activation of *Nmbr*- or *Grpr* -expressing preBötC neurons induces sighs

Activation of *Nmb*- or *Grp*-expressing pF neurons generates sufficient input to preBötC to generate ectopic sighs, but is activation of neuropeptide-receptor expressing preBötC neurons via their cognate receptors necessary for sigh production? We tested whether bypassing the peptidergic receptors and directly activating preBötC *Grpr*- or *Nmbr*-expressing neurons could also generate sighs. To do this, we expressed ChR2 or the excitatory DREADD hM3Dq in discrete subsets of preBötC neurons (see *Methods*).

To express ChR2 in preBötC GRPR neurons, we microinjected a Flp-dependent ChR2 virus^10^ into the preBötC of Grpr^Flp^ mice (Extended data Fig. 2c). While no ectopic sighs could be generated by bilateral preBötC SPP in these mice (phase: 0.0-1.0), preBötC LPP during 0.2–0.9 sigh phases elicited ectopic sighs which appeared as partial doublet or augmented breaths, and reset the sigh cycle (Fig. 3a, bottom). Ectopic sighs elicited by preBötC LPP of GRPR neurons were no different from spontaneous sighs in V_T_ and *T_E_* (Fig. 3b) and exhibited a postsigh apnea (*T_E_* following evoked sighs increased to 194 ± 22%, Fig. 3b).

**Fig. 3.**
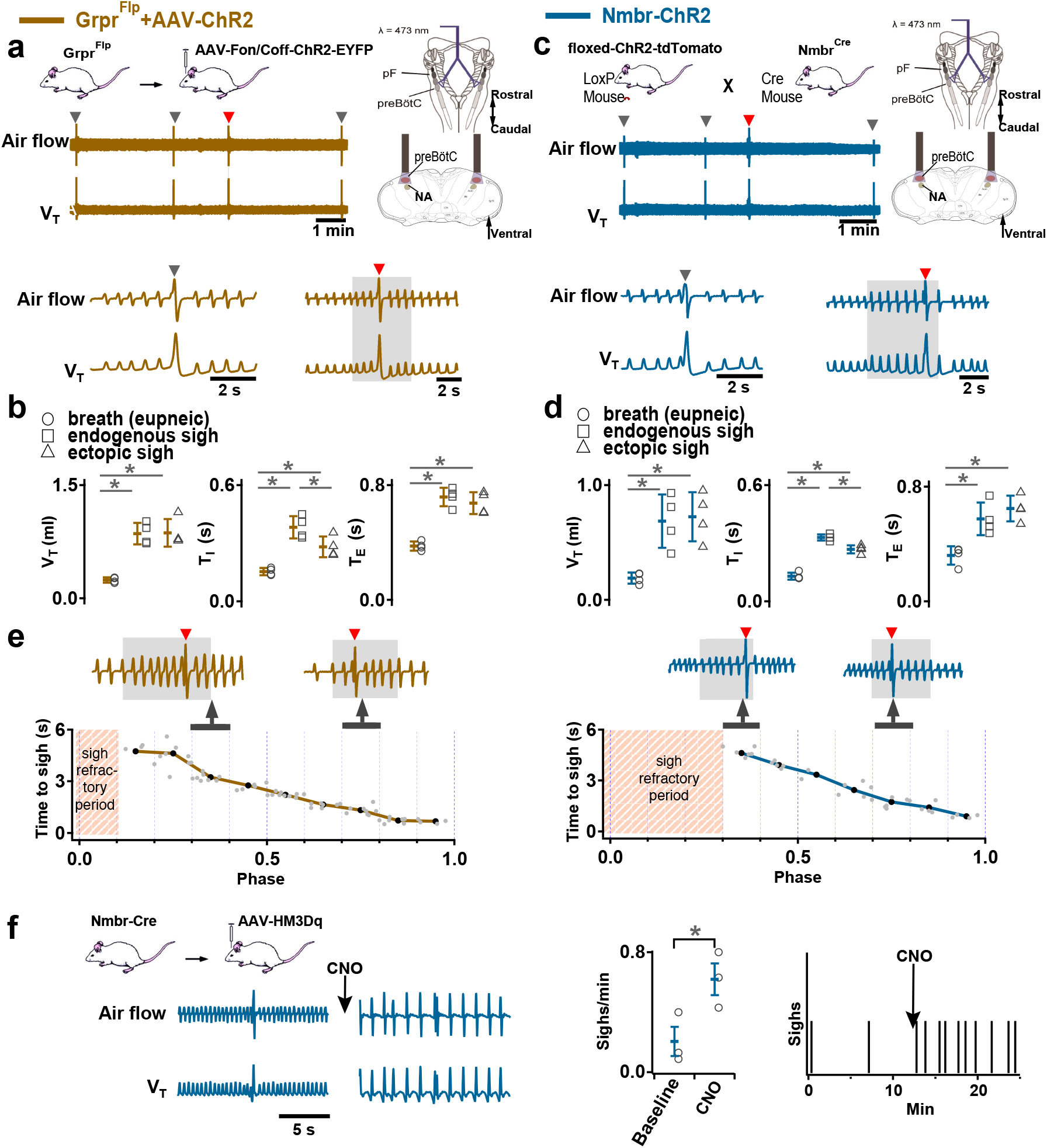
Excitation of preBötC GRPR or NMBR neurons induced sighs. (a-e) Photostimulation of preBötC GRPR (brown) or NMBR (blue) neurons evoked sighs. **a**, Top: Schematic of genetic strategy to target preBötC GRPR neurons. Schematic (right) depicting bilateral placement of optical cannula targeting preBötC. Middle and Bottom: Raw (middle) and expanded (bottom) traces show bilateral LPP (gray box) of preBötC GRPR neurons could elicit an ectopic sigh (red arrowhead). Gray arrowheads indicate endogenous sighs. **b**, Photo- stimulation of preBötC GRPR induced ectopic sighs with similar V_T_ and *T_E_* as endogenous sighs, but slightly lower *T_I_* (V_T_: t_3_ = 0.233, p = 0.824; *T_E_*: t_3_ = 0.249, p = 0.812; *T_I_*: t_3_ = 3.267, p < 0.05). V_T_, *T_E_* and *T_I_* of ectopic sighs were significantly greater than those of eupneic breaths (V_T_: t_3_ = 6.889, p = 5 x 10^-4^; *T_I_*: t_3_ = 3.183, p = 0.019; *T_E_*: t_3_ = 8.675, p = 3 x 10^-4^). Statistical significance was determined with a One Way RM ANOVA followed by All Pairwise Multiple Comparison Procedures (Holm-Sidak method), V_T_: F_2,9_ = 32.744, p < 0.001; *T_I_*: F_2,9_ = 16.530, p < 0.001; *T_E_*: F_2,9_ = 48.771, p < 0.001. **c**, Top: Targeting scheme to generate Nmbr-ChR2 mice. Schematic (right) depicting bilateral placement of optical cannula targeting preBötC. Middle and Bottom: Raw (middle) and expanded (bottom) show bilateral LPP (gray box) of preBötC NMBR neurons could elicit an ectopic sigh (red arrowhead). Gray arrowheads indicate endogenous sighs. **d**, Photostimulation of preBötC NMBR neurons induced ectopic sighs with similar V_T_ and *T_E_* as endogenous sighs, but slightly lower *T_I_* (V_T_: t_3_ = 0.820, p = 0.444; *T_E_*: t_3_ = 0.976, p = 0.367; *T_I_*: t_3_ = 3.175, p = 0.0192). V_T_, *T_E_* and *T_I_* of ectopic sighs were significantly greater than those of eupneic breaths (V_T_: t_3_ = 9.756, p = 7 x 10^-5^; *T_I_*: t_3_ = 7.685, p = 3 x 10^-4^; *T_E_*: t_3_ = 7.326, p = 3 x 10^-4^). Statistical significance was determined with a One Way RM ANOVA followed by All Pairwise Multiple Comparison Procedures (Holm-Sidak method), V_T_: F_2,9_ = 58.564, p < 0.001; *T_I_*: F_2,9_ = 62.362, p < 0.001; *T_E_*: F_2,9_ = 31.646, p < 0.001. **e**, Latency from laser onset was negatively correlated with phase in Grpr^Flp^ (r = -0.980, p = 4 x 10^-6^) and Nmbr-ChR2 (r = -0.994, p = 6 x 10^-6^) mice (Pearson Product Moment Correlation). Representative traces showing latency between laser onset (gray box indicates LPP) and ectopic sighs (red arrowheads) when LPP was applied in medial (0.4) and late (0.8) phase. Gray dots represent latency to ectopic sigh from stimulation onset in each phase in Grpr^Flp^ (n = 5 mice) and Nmbr-ChR2 (n = 4 mice) mice, solid lines join the average latency in each phase (black circle). No sighs were generated by LPP in phase 0.0–0.1 (23 ± 4 s from previous sigh, measured from 55 stimulus in 5 mice: basal sigh rate 16 ± 2 h/hour, range 12–19; intersigh interval 230 ± 41 s) in Grpr^Flp^ and in phase 0.0–0.3 (61 ± 21 s from previous sigh, measured from 50 stimulus in 4 mice: basal sigh rate 19 ± 6 h/hour, range 12-25; intersigh interval 203 ± 70 s) in Nmbr-ChR2 mice. **f**, Chemogenetic activation of preBötC NMBR neurons induced sighs. Left top: schematic diagram of genetic strategy to selectively express DREADD receptor hM3Dq on preBötC NMBR neurons. Left bottom: representative traces of airflow and V_T_ before and after application of CNO to brainstem surface. Middle: Activation of hM3Dq receptors expressed on NMBR neurons with CNO significantly increases sigh rate (paired two-tailed t-test, n = 3 mice: t_2_ = 5.94, p = 0.03). Right: Trace from a representative mouse illustrating incidence of sighs before and after CNO application. Data are shown as mean ± SE. Asterisks indicate post-hoc multiple comparison test results or paired t- test results: *, significance with p < 0.05.

Similarly, in Nmbr-ChR2 mice (Fig. 3c, top), preBötC SPP (phase: 0.0-1.0) had no effect on sighing, while LPP during 0.4-0.9 sigh phases elicited ectopic sighs which appeared as a partial doublet or an augmented breath, and reset the sigh cycle (Fig. 3c, bottom). V_T_ and postsigh *T_E_* of ectopic sighs were no different from that of spontaneous sighs (Fig. 3d). Ectopic sighs elicited by preBötC LPP of NMBR neurons exhibited a postsigh apnea (postsigh *T_E_* increased to 225 ± 33%, Fig. 3d).

As with photostimulation of pF NMB and GRP neurons, there was a postsigh refractory period during which we could not generate ectopic sighs in preBötC NMBR or GRPR neurons. Again, the latency from LPP to ectopic sigh onset was inversely related to phase (Fig 3e).

We also tested whether activation of preBötC NMBR neurons using an excitatory DREADD could induce sighing. To do this, we transduced preBötC *Nmbr*-expressing neurons by microinjection of Cre-dependent AAV-hM3Dq in Nmbr^Cre^ mice. In these mice, application of the selective agonist clozapine-n-oxide (CNO) to the brainstem surface increased sigh frequency ∼threefold (Fig. 3f; see Extended data Fig.4 for control CNO application). Sighs induced by CNO in Nmbr^Cre^ mice were of the doublet shape. Thus, increasing the excitability of preBötC NMBR neurons increases sighing.

**Fig. 4.**
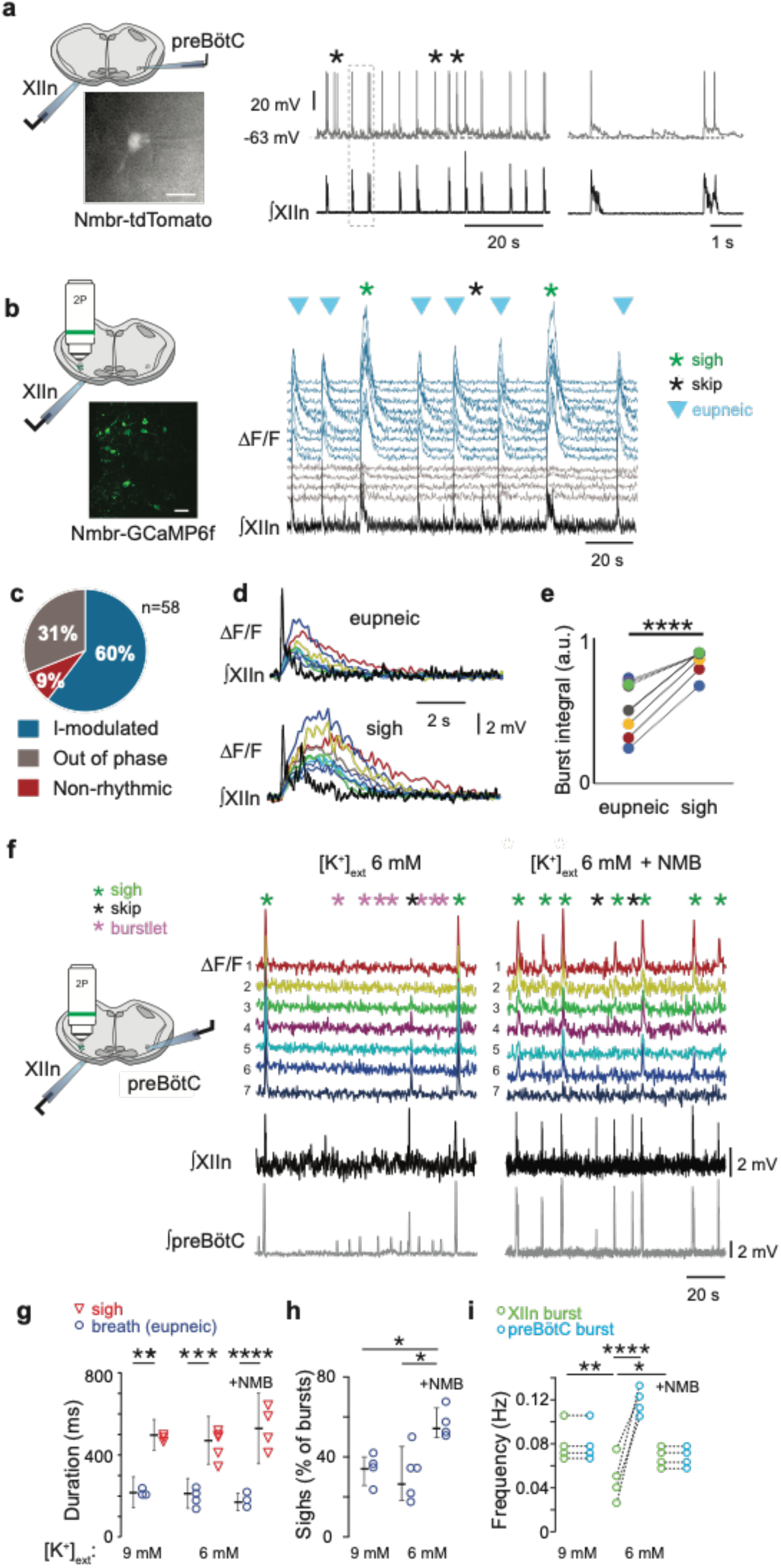
NMBR neurons in preBötC were rhythmically active *in vitro* and NMB converted inspiratory burstlets into sighs and bursts. **a**, Left: schematic of slice preparation with photograph of patched Nmbr-tdTomato preBötC neuron. Scale bar: 50 µm. Right: Whole cell current clamp recording from inspiratory-modulated preBötC NMBR neuron (top trace_ that generated action potentials (APs) during each XII burst (bottom trace). Not every cluster of APs (*) is associated with a burst, but each burst was associated with one or more Aps. Far right: Two bursts shown at expanded timescale. **b**, Left: schematic of slice preparation in 2P experiment with image of GCaMP6f-expressing preBötC neurons. Scale bar 50 µm. Right: Representative Ca^2+^ signal of neighboring preBötC NMBR neurons was correlated with XIIn bursts (9 mM [K^+^]_ext_). Large amplitude XIIn bursts represent sighs (green asterisks), XIIn bursts not associated with Ca^2+^ signal indicated with black asterisks. **c**, Percentage of rhythmically active (I-modulated), non-rhythmic and out of phase preBötC NMBR neurons in preBötC. **d**, Ca^2+^ transients in individual neurons were significantly larger during sighs than during eupneic bursts (paired two-tailed t-test: t_8_ = 7.14, p < 0.001). **e**, Intervals of sighs were significantly greater during sighs compared to eupneic burst (same color-coded neurons as in d). **f**, Left: schematic of slice preparation and configuration. Right: Simultaneous Ca2+ signal from NMBR neurons (n = 7), preBötC field recording, and XIIn activity in control conditions ([K^+^]_ext_ = 6 mM) and after addition of NMB (30 nM). Burstlets (red asterisks) are seen in control but not with NMB. **g**, Quantification of duration of eupneic and sigh bursts in 9mM [K^+^]_ext_ and 6mM [K^+^]_ext_ with or without NMB. Differences in duration between eupneic bursts in different conditions, and between sighs in different conditions were not significant (repeated measures ANOVA, F_5,19_ = 20.1, p < 0.001; asterisks indicate significance from Tukey post-hoc tests). **h**, Percentage of bursts that were sighs during 5 min was significantly higher in 6 mM [K^+^]_ext_ + NMB than other conditions (repeated measures ANOVA, F_2,9_ = 7.18, p = 0.018). **i**, Comparisons between XIIn burst frequency and preBötC burst frequency in in 9mM [K^+^]_ext_ and 6mM [K^+^]_ext_ with or without NMB. XIIn and preBötC burst frequency were similar in 9mM [K^+^]_ext_ and 6mM [K^+^]_ext_ + NMB, and both conditions differ from 6mM [K^+^]_ext_ alone. There is an increase in preBötC burst frequency in 6mM [K+]ext due to the burstlets (shown in a), (repeated measures ANOVA, F_5,23_ = 11.57, p < 0.001). Data are shown as mean ± SE. Asterisks indicate significance from Tukey post-hoc tests or paired t-tests: *, significance with p < 0.05; **, significance with p < 0.01; ***, significance with p < 0.001.

### preBötC NMBR neurons are rhythmically active

We next asked: what are the electrophysiological properties of the *Grpr*- and *Nmbr*-expressing preBötC neurons? We took several approaches. First, using inspiratory rhythmic slices^11^ from neonatal Grpr-EGFP or Nmbr-tdTomato reporter mice, we obtained whole-cell patch-clamp recordings from preBötC GRPR or NMBR neurons while monitoring hypoglossal nerve (XII) inspiratory activity. In Grpr-EGFP mice, only 3 of 21 preBötC GRPR neurons were inspiratory-modulated, i.e., fired during XII inspiratory bursts. In contrast, in Nmbr-tdTomato mice, the majority of preBötC NMBR neurons (10/16) were inspiratory-modulated (Fig. 4a). This finding was confirmed using two-photon calcium imaging of rhythmic slices from Nmbr-GCaMP6f mice, in which we could monitor the activity of up to 10 neurons simultaneously (Fig. 4b). The majority of preBötC NMBR neurons (35/58; n = 6) exhibited inspiratory-modulated Ca^2+^ oscillations coincident with XII bursts (Fig. 4b); however, episodes of non-breathing-modulated episodic bursting, i.e., bursts not in phase with XII, were also seen (Fig. 4c) and varied in frequency from 0.025 to 0.1 Hz (Extended Data Fig. 5). Almost all NMBR neurons that had Ca^2+^ elevations during inspiratory bursts also had elevations during sighs (34/35), but not during the pre-I period, consistent with whole-cell recordings (Fig. 4a), indicating that they are likely Type II, i.e., presumptive non-rhythm generating, neurons^12^ (Fig. 4d). The burst duration of the GCaMP6f signal associated with sighs was longer-lasting than those associated with eupneic events in the same neuron (Fig. 4d-e).

### NMB converts NMBR neurons inspiratory burstlets into sighs and bursts

In typical recording conditions *in vitro* ([K^+^]_ACSF_ = 9 mM), the preBötC population activity burst and the periodic XII discharges are in sync. Under conditions of low excitability ([K^+^]_ACSF_ < 7 mM), however, in addition to these coincident high amplitude bursts in preBötC and XII, there are low amplitude burstlets in preBötC but not in XII^11^. We have hypothesized that the burstlets represent the rhythmogenic kernel^11,13,14^. NMBR neurons had a Type II firing pattern, i.e., no preinspiratory spiking, suggesting that they are not rhythmogenic.

We reliably observed both burstlets and bursts in population activity when we lowered [K^+^]_ACSF_ to 6 mM^9^. When we did so, NMBR neurons were still active during bursts but never during burstlets (Fig. 4f). When we applied NMB (30 nM), only bursts were observed, while both burst and sigh frequency in both preBötC and XII increased (Fig. 4g). The duration of sighs and eupneic bursts were similar in [K^+^]_ACSF_ = 6 mM vs 9 mM (Fig. 4h), while the percentage of sighs increased by 30% during NMB (30 nM) application in 6 mM K^+^ (Fig. 4i), similar to the increase elicited by 30 nM NMB at 9 mM K^+^ ^4^.

### Activating preBötC GRPR- or NMBR-only neurons has distinct effects on sighing

Photoactivation of preBötC GRPR or NMBR neurons can induce sighs. Given that there are three bombesin-related peptide receptor-expressing subpopulations in preBötC, i.e., neurons that express *Nmbr* but not *Grpr* (NMBR-only), neurons that express *Grpr* but not *Nmbr* (GRPR- only) and neurons that express both *Nmbr* and *Grpr*, does activation of neurons expressing one but not the other of these receptors affect sighs differently? We used an intersectional genetic approach to target neurons that express only one of these receptors, excluding double-positive (GRPR/NMBR) neurons^15^. Thus, using Cre-on/Flp-off and Cre-off/Flp-on viral vectors for the delivery of ChR2 in Nmbr^Cre^;Grpr^Flp^ double mutant mice, we expressed ChR2 in preBötC NMBR- only (Fig. 5a) or GRPR-only neurons (Fig. 5d), respectively. We examined the effect of preBötC LPP and SPP of these neurons on sighing. When targeting NMBR-only neurons, preBötC SPP elicited ectopic bursts which appeared similar to partial doublets or augmented breaths with smaller V_T_ compared to spontaneous sighs due to a decrease in *T_I_* to 64 ± 8% (Fig. 5c). *T_E_* immediately following these ectopic bursts were longer than the *T_E_* of eupneic breaths (Fig. 5c).

**Fig. 5.**
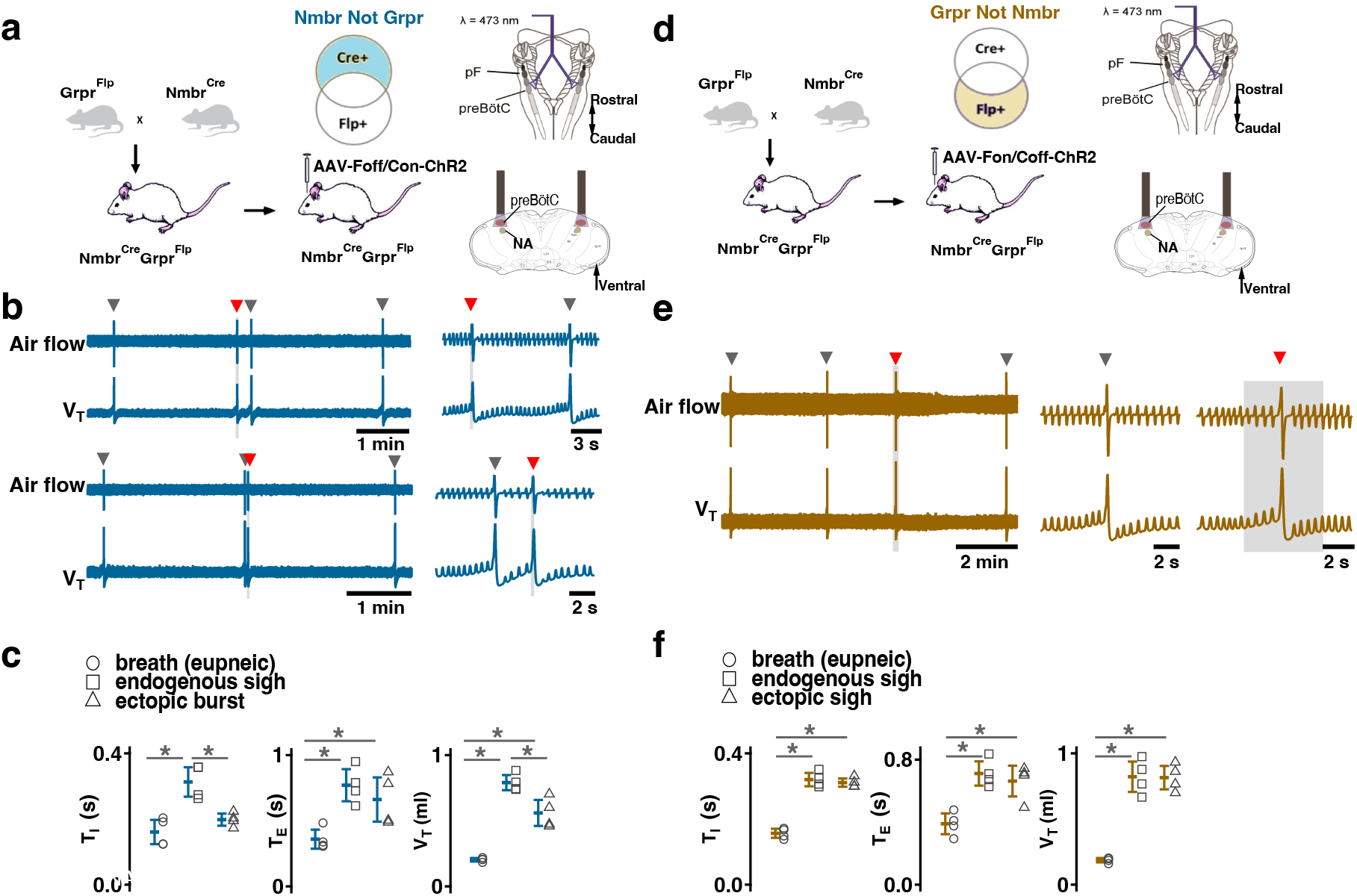
Effects of activating preBötC NMBR-only or GRPR-only neurons on the generation of sighs. (a-c) Effects of activating preBötC NMBR-only neurons on sigh generation. **a,** Left: Schematic of the intersectional genetic strategy to target preBötC NMBR-only neurons. Right: Schematic depicting bilateral placement of optical cannula targeting preBötC. **b,** Top: Raw (left) and expanded (right) traces show that preBötC SPP (gray box) of NMBR-only neurons induced an ectopic burst (red arrowhead) which did not reset the sigh rhythm. Bottom: preBötC SPP of NMBR-only neurons induced an ectopic burst (red arrowhead) in a sigh refractory period. Gray arrowheads indicate endogenous sighs. **c,** V_T_ of ectopic bursts evoked by NMBR-only photostimulation was smaller than that of endogenous sighs (n = 4 mice, t_3_ = 4.278, p = 0.005), but larger than of eupneic breaths (t_3_ = 6.588, p = 6 x 10^-4^); *T_E_* of evoked ectopic bursts was no different from endogenous sighs (t_3_ = 1.882, p = 0.109), but higher than of eupneic breaths (t_3_ = 5.203, p = 0.002); *T_I_* of evoked ectopic bursts was shorter than that of endogenous sighs (t_3_ = 3.917, p = 0.008). Statistical significance was determined with a One Way RM ANOVA followed by All Pairwise Multiple Comparison Procedures (Holm-Sidak method), V_T_: F_2,9_ = 59.922, p < 0.001; *T_I_*: F_2,9_ = 25.206, p < 0.001; *T_E_*: F_2,9_ = 26.939, p < 0.001. (d-f) Effects of activating preBötC GRPR-only neurons on sigh generation. **d,** Left: Schematic of the intersectional genetic strategy to target preBötC GRPR-only neurons. Right: Schematic depicting bilateral placement of optical cannula targeting preBötC. **e,** preBötC LPP of GRPR- only neurons in late phase elicited ectopic sighs and reset the sigh rhythm. Raw (left) and expanded (middle and right) show bilateral LPP (gray box) of preBötC GRPR-only neurons elicited an ectopic sigh (red arrowhead). Gray arrowheads indicate endogenous sighs. **f,** V_T_ (t_3_ = 0.152, p = 0.884), *T_I_* (t_3_ = 1.843, p = 0.115) and *T_E_* (t_3_ = 1.619, p = 0.157), of ectopic sighs were no different from endogenous sighs (n = 4 mice), but increased compared to eupneic breaths (V_T_:, t_3_ = 12.855, p = 10^-5^; *T_I_*: t_3_ = 30.235, p = 9 x 10^-8^; *T_E_*: t_3_ = 9.007, p = 4 x 10^-5^), indicative of augmented breaths with postsigh apneas. Statistical significance was determined with a One Way RM ANOVA followed by All Pairwise Multiple Comparison Procedures (Holm- Sidak method), V_T_: F_2,9_ = 111.488, p < 0.001; *T_I_*: F_2,9_ = 648.855, p < 0.001; *T_E_*: F_2,9_ = 65.544, p < 0.001. Data are shown as mean ± SE. Asterisks indicate post-hoc multiple comparison test or paired t-test results: *, significance with p <0 .05.

Note that the time between the SPP-induced ectopic burst and the next endogenous sigh was shorter than the time between endogenous sighs, indicating that SPP-induced burst did not reset the sigh cycle (Fig. 5b, top); SPP-induced ectopic burst could also occur in a sigh refractory period right after the previous eupneic sigh (Fig. 5b, bottom). When targeting GRPR- only neurons, while ectopic sighs could not be generated by SPP at any phase of the sigh cycle, preBötC LPP in the late phase of the sigh cycle elicited ectopic sighs with a partial doublet or augmented breath shape and reset the sigh cycle (Fig. 5e). *T_I_*, *T_E_* and V_T_ of LPP-induced sighs were indistinguishable from spontaneous sighs (Fig. 5f). The ectopic sighs exhibited a postsigh apnea, *T_E_* after evoked sighs increased to 171 ± 12% of the *T_E_* of eupneic breaths (Fig. 5f). These data suggest that GRPR and NMBR neurons play different overlapping roles in sigh generation.

### Activating preBötC GRPR or NMBR neurons has distinct effects on inspiratory burst amplitude

In addition to generating sighs, activation of preBötC NMBR and/or GRPR neurons also affected eupneic breathing frequency (*f*) and V_T_. In Grpr^Flp^ mice expressing ChR2 in preBötC, LPP during the sigh refractory period did not generate ectopic sighs but increased *f* to 131 ± 9% of baseline (Fig. 6a), and increased V_T_ to 117 ± 4% of baseline (Fig. 6a), however, when excluding double-positive (GRPR/NMBR) neurons, preBötC SPP or LPP of GRPR-only neurons did not change the inspiratory burst amplitude in stimulus cycles (Fig. 6b).

**Fig. 6.**
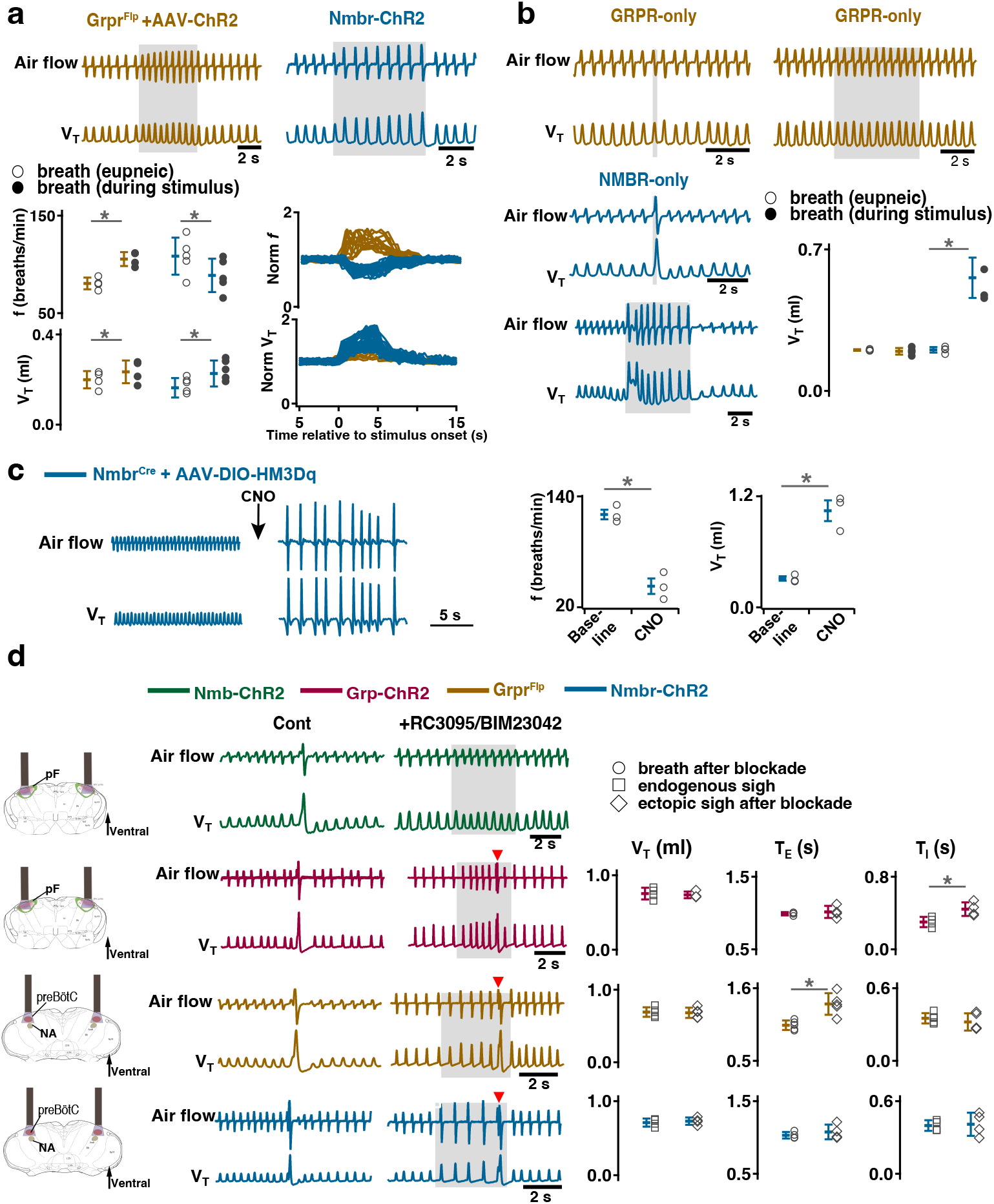
Distinct roles of preBötC GRPR and NMBR neurons on inspiratory burst amplitude and sigh generation after receptor blockade. a, Photostimulation of preBötC GRPR (brown) and NMBR (blue) neurons perturbs breathing frequency and inspiratory burst amplitude. Top: representative traces recorded during sigh refractory period show effects of bilateral preBötC LPP of preBötC GRPR (brown) or NMBR (blue) neurons on eupneic breathing f and V_T_. Bottom: V_T_ and *f* of eupneic breaths and during photostimulation. (paired t-tests, Grpr^Flp^ + AAV-ChR2, n = 4 mice*, f*: t_3_ = -8.13801, p = 4 x 10^-3^; V_T_: t_3_ = -5.035, p = 0.015; Nmbr-ChR2, n = 5 mice, *f*: t_4_ = 7.33379, p = 2 x 10^-3^, V_T_: t_4_ = -5.875, p = 4 x 10^-3^). Normalized (norm) V_T_ and *f* traces relative to stimulus onset measured prestimulation, during stimulation and poststimulation. **b,** Distinct effects of activating preBötC GRPR-only (brown) and NMBR-only (blue) neurons on inspiratory burst amplitude. Top and bottom (left): representative traces show the effects of bilateral preBötC SPP and LPP of preBötC GRPR-only (brown) or NMBR-only (blue) neurons on breathing V_T_. Bottom (right): V_T_ of eupneic breaths and during SPP stimulation. Increase in V_T_ in response to bilateral preBötC SPP of preBötC GRPR-only (brown) or NMBR-only (blue) neurons (paired t-tests, GRPR-only, n = 4 mice: t_3_ = 0.746, p = 0.510; NMBR-only, n = 4 mice: t_3_ = -6.886, p = 6 x 10^-3^). **c,** Activation of hM3Dq receptors expressed on NMBR neurons with CNO significantly increased breathing amplitude and decreased *f* (paired two-tailed t-tests, n = 3 mice, *f*: t_2_ = 5.71, p = 0.03; V_T_: t_2_ = 5.47, p = 0.03). **d,** Sighs can be induced after GRPR and NMBR inhibition. Left: schematic depicting bilateral placement of optical cannula targeting pF or preBötC. Middle: representative traces showing LPP of pF GRP neurons, or preBötC GRPR and NMBR neurons elicited ectopic sighs (doublet, red arrowhead), but not of pF NMB neurons, in the presence of GRPR antagonist RC3095 and NMBR antagonist BIM23042. Right: V_T_ of doublets induced by pF LPP of GRP neurons was not significantly different from that of endogenous sighs (n = 4 mice; t_3_ = 0.724, p = 0.522); *T_I_* increased compared to endogenous sighs (t_3_ = -7.493, p = 5 x 10^-3^), *T_E_* of the following respiratory cycle was unaffected (t_3_ = -0.472, p = 0.669). V_T_ and *T_I_* of the doublets elicited by preBötC LPP of GRPR (n = 5 mice, V_T_: t_4_ = 0.199, p = 0.852; *T_I_*: t_4_ = 0.912, p = 0.413) or NMBR (n = 4 mice, V_T_: t_3_ = -0.845, p = 0.460; *T_I_*: t_3_ = -0.376, p = 0.732) neurons after blockade of both receptors were not different from spontaneous sighs. *T_E_* immediately following doublet increased in Grpr^Flp^ (n = 5 mice, t_4_ = -3.144, p = 0.035), but not lengthened in Nmbr-ChR2 mice (n = 4 mice, t_3_ = -0.801, p = 0.482). Data are shown as mean ± SE. Asterisks indicate post-hoc multiple comparison test or paired t-test results: *, significance with p < 0.05.

In contrast, in Nmbr-ChR2 mice, preBötC LPP delivered during a postsigh refractory period did not generate ectopic sighs but decreased *f* to 83 ± 4% (Fig. 6a) and increased V_T_ to 140 ± 12% (Fig. 6a) of baseline. preBötC SPP or LPP of NMBR-only neurons increased V_T_ to 274 ± 41% or 264 ± 17% of control, respectively (Fig. 6b).

The effect on inspiratory burst amplitude was also observed in Nmbr^Cre^ mice that had preBötC NMBR neurons transfected with Cre-dependent AAV-HM3Dq; CNO applied to the brainstem surface significantly increased V_T_ by 343 ± 56% (Fig. 6c) and decreased *f* by 63 ± 8.4% (Fig. 6c). Note that although such breaths were of the same V_T_ as eupneic sighs before administration of CNO (Fig.3c), in line with our definition of sighs (Extended Data Fig.1), we classify this outcome as large amplitude breaths rather than “all sigh” breathing rhythm, according to our definition of sighs (Extended Data Fig.1).

Thus, stimulation of preBötC NMBR and/or GRPR neurons also has a profound effect on the eupneic, nonsigh, breathing pattern, and GRPR and NMBR neurons appear to differently affect inspiratory burst amplitude.

### Sighs can be induced after inhibition of GRPRs or NMBRs

Expression of *Grpr* and/or *Nmbr* defines subsets of preBötC neurons that affect sigh generation. Is activation of these peptide receptors *necessary* for sigh generation?

For that we investigated the effects of activating peptide-expressing pF neurons in the presence of bombesin receptor antagonists. First, we confirmed that bilateral microinjection of the NMBR antagonist BIM23042 and GRPR antagonist RC3095 (300 µM each, 50 nl/side) into the preBötC in anesthetized mice eliminates spontaneous sighs (Extended Data Fig. 6). Next, we determined the effects of such NMBR and GRPR antagonism on LPP-evoked sighs. After blockade, no sighs were elicited by pF LPP in Nmb-ChR2 mice; in contrast, pF LPP of GRP neurons still elicited ectopic sighs of a doublet shape. The V_T_ was not significantly different from that of spontaneous sighs; *T_E_* of the following respiratory cycle was unaffected (Fig. 6d). Thus, when pF NMB neurons were stimulated, preBötC NMBR neurons require NMBR activation to contribute to sigh generation, whereas when pF GRP neurons were stimulated, sighs could still be generated after antagonism of preBötC GRPR receptors.

We next investigated whether explicit activation of preBötC NMBRs and/or GRPRs is necessary to produce sighs, i.e., could sighs be generated simply by depolarizing these preBötC neurons to fire action potentials? In ChR2-transduced Grpr^Flp^ or Nmbr-ChR2 mice after antagonism of both receptors sufficient to eliminate spontaneous sighs (Extended Data Fig. 6), LPP of preBötC GRPR or NMBR neurons still elicited sighs of doublet shape. V_T_ and *T_I_* of the sighs elicited in Grpr^Flp^ or Nmbr-ChR2 mice after blockade of both receptors was no different from spontaneous sighs (Fig. 6d). Postsigh apnea was observed in Grpr^Flp^ (*T_E_* immediately following the sigh increased to 132 ± 24%), but not in Nmbr-ChR2 mice (Fig. 6d).

Thus, initiating or increasing the activity of pF GRP neurons, or preBötC GRPR or NMBR neurons, can generate sighs without activation of these receptors, either endogenously or exogenously.

### preBötC SST neurons mediate sighing

Can activity of preBötC neurons other than those expressing *Grpr* or *Nmbr* generate sighs? To address this question, we virally expressed ChR2 (Fig. 7a), hM3Dq (Fig. 7e), or the PSAM4 (Fig. 7g) in preBötC SST neurons. In ChR2-transduced Sst^Cre^ mice, bilateral preBötC SPP during inspiration elicited ectopic sighs of partial doublets or augmented breath shapes and reset the sigh cycle (Fig. 7b). V_T_, *T_I_* and *T_E_* of the induced sighs were no different from those of spontaneous sighs (Fig. 7c). The ectopic sighs elicited by photostimulation of preBötC SST neurons exhibited a postsigh apnea, *T_E_* of the following respiratory cycle increased by 210 ± 19% (Fig. 7c). After BIM23042 and RC3095 microinjection (300 µM each, 50 nl/side), spontaneous sighs were eliminated (Extended Data Fig. 6). In the presence of antagonists, preBötC SPP during inspiration could still elicit an ectopic sigh (Fig. 7d). V_T_ and *T_I_* of ectopic sighs elicited after blockade of both receptors were no different from spontaneous sighs, or from ectopic sighs elicited before blockade (Fig. 7c). Ectopic sighs elicited after blockade also had a postsigh apnea, *T_E_* of the following respiratory cycle increased by 236 ± 49% (Fig. 7c).

**Fig. 7.**
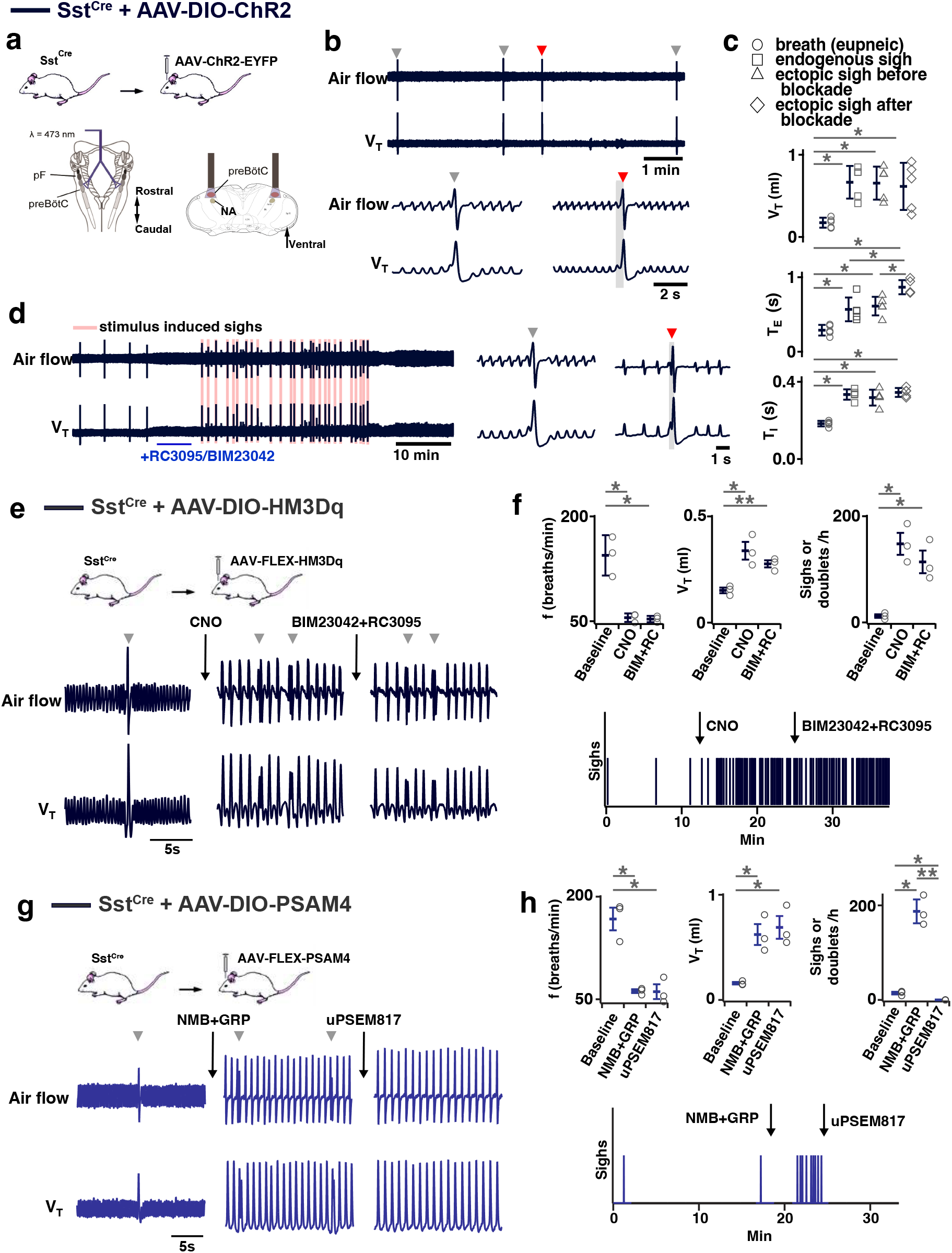
**Effects of activation or inhibition of preBötC SST neurons on sigh generation**. (a-d) Optogenetic activation of SST neurons generates sighs after blockade of NMBRs and GRPRs. **a,** Top: Schematic of the genetic strategy to target preBötC SST neurons. Bottom: Schematic depicting bilateral placement of optical cannula targeting preBötC. **b,** Ectopic sigh (red arrowhead) elicited by bilateral SPP (gray box) of preBötC SST neurons. **c,** V_T_ (t_4_ = 0.176, p = 0.863), *T_I_* (t_4_ = 1.355, p = 0.200), *T_E_* (t_4_ = 0.734, p = 0.477) of ectopic sighs before blockade or V_T_ (t_4_ = 0.856, p = 0.409), *T_I_* (t_4_ = 0.894, p = 0.389) of ectopic sighs after blockade of NMBR and GRPR were no different from endogenous sighs (n = 5 mice). *T_E_* of ectopic sighs elicited after blockade was longer than that of endogenous sighs (t_4_ = 4.960, p = 3 x 10^-4^). V_T_, *T_I_*, and *T_E_* of ectopic sighs before; V_T_: t_4_ = 7.925, p = 4 x 10^-6^; *T_I_*: t_4_ = 11.194, p = 10^-7^; *T_E_*: t_4_ = 5.405, p = 10^-4^) or after (V_T_: t_4_ = 7.245, p = 10^-5^; *T_I_*: t_4_ = 13.442, p = 10^-8^; *T_E_*: t_4_ = 9.631, p = 5 x 10^-7^) blockade were increased compared with eupneic breaths (n = 5 mice), indicative of augmented breaths with postsigh apneas. Statistical significance was determined with a One Way RM ANOVA followed by All Pairwise Multiple Comparison Procedures (Holm-Sidak method), V_T_: F_3,16_ = 30.357, p < 0.001; *T_I_*: F_3,16_ = 78.526, p < 0.001; *T_E_*: F_3,16_ = 31.134, p < 0.001. **d,** SPP elicits sighs in the presence of GRPR antagonist RC3095 and NMBR antagonist BIM23042. (e- f) Chemogenetic activation of SST neurons generated sighs after blockade of NMBRs and GRPRs. **e,** Top: schematic diagram of genetic strategy to selectively express DREADD receptor hM3Dq on preBötC SST neurons. Bottom: representative trace of airflow and V_T_ during baseline, after application of CNO, and after microinjection of RC3095 and BIM23042. **f,** Top: Activation of hM3Dq receptors expressed on preBötC SST^+^ neurons significantly decreased breathing *f*, increases V_T_, and elevated sigh frequency; subsequent BIM23042 and RC3095 (B+R) microinjection into preBötC did not significantly affect breathing *f*, V_T_, nor sigh rate induced by CNO application (repeated measures ANOVA, n = 3 mice; *f*: F_2,4_ = 28.3, p = 0.004; V_T_: F_2,4_ = 25.7, p = 0.005; sigh rate: F_2,4_ = 26.1, p = 0.005). Bottom: representative trace depicting sighs during baseline, after application of CNO and after microinjection of NMBR and GRPR antagonists RC3095 and BIM23042. **g,** Top: schematic diagram of genetic strategy to selectively express ultrapotent inhibitory DREADD receptor PSAM4-GlyR on preBötC SST^+^ neurons. Bottom: representative trace of airflow and V_T_ during baseline, after preBötC microinjection of peptides NMB and GRP (250 µM each, 50nl/side), and application of uPSEM817 (10mM, 30µl applied to brainstem surface). **h,** Top: Microinjection of peptides NMB and GRP into preBötC significantly decreased breathing *f,* increased V_T_, and elevated sigh frequency; subsequent inhibition of SST+ preBötC neurons selectively eliminated any sighs, but preserves decreased *f* and V_T_ (repeated measures ANOVA, n = 3 mice; f: F_2,4_ = 55.8, p = 0.001; V_T_: F_2,4_ = 19.0, p = 0.009; sigh rate: F_2,4_ = 83.8, p < 0.001). Bottom: representative trace depicting sighs during baseline, after bilateral preBötC microinjection of NMB and GRP and after application of PSAM4-GlyR ligand uPSEM817 to the brainstem surface. Data are shown as mean ± SE. Asterisks indicate post-hoc multiple comparison test or paired t-test results: *, significance with p < 0.05; **, significance with p < 0.01.

In hM3Dq-transduced Sst^Cre^ anesthetized mice (Fig. 7e), excitation of preBötC SST neurons by CNO application onto ventral brainstem surface affected eupneic breathing: *f* decreased by 60 ± 6.2%, V_T_ increased by 222 ± 21%. CNO application increased sigh rate ∼tenfold with sighs appearing as doublets (Fig. 7f). In these mice, subsequent microinjection of BIM23042 and RC3095 into preBötC did not affect *f*, V_T_ or sigh rate induced by CNO administration (Fig. 7f). Thus, excitation of preBötC SST neurons can produce sighs in an NMBR- and GRPR- independent manner.

Next, we investigated whether sighs induced by the NMBR- and GRPR-dependent pathway requires activity of preBötC SST neurons. For that we transfected preBötC SST neurons with Cre-dependent ultrapotent inhibitory DREADD AAV-PSAM4-GlyR^16^ in Sst^Cre^ mice (Fig. 7g). The sigh rate was then increased by microinjection of peptides NMB and GRP (250 µM each, 50 nl/side) into preBötC (Fig.7g, h) in ketamine/xylazine anesthestized mice, which also decreased *f*, and increased V_T_ (Fig.7g, h). Subsequent application of the PSAM4-GlyR ligand uPSEM817 to the brainstem surface did not affect *f* or V_T_, but completely eliminated all sighs (Fig.7g, h).

Note that similar to excitatory DREADD effects in Nmbr^Cre^ mice, manipulations increasing V_T_ of all breaths were classified to affect V_T_ of eupneic breaths rather than induce an “all sigh” rhythm, according to our definition of sighs (Extended Data Fig.1).

## Discussion

In all mammals, sighs are generated periodically to maintain lung function and, in humans, are episodically associated with such emotional states as relief, sadness, yearning, exhaustion, stress, and joy^1–3,6,7^, and in rodents are associated with inferred indicators of emotional states such as stress and anxiety^17^. The signals producing these large periodic inspiratory efforts originate within the preBötC, a heterogeneous medullary population that is the kernel for generation of breathing rhythm^18^. Molecularly-defined preBötC subpopulations underlie different functions during breathing^18^. Ablation, genetic deletion or pharmacological excitation or inhibition of preBötC neurons expressing receptors (NMBR and GRPR) for bombesin-related peptides (NMB and GRP) significantly affect rhythmic (∼25-40/hour) sighing in rodents^4^. Here, we further explored the mechanisms of sigh generation.

Sighs exhibit various shapes, i.e., augmented single inspiratory effort, partial doublets, and full doublets, that have two unifying characteristics: i) significant transient increase in V_T_ and; ii) occur at a much lower frequency compared to eupneic breaths; often these augmented inspiratory efforts are followed by a lengthened interburst interval, i.e., postsigh apnea. We use these criteria as a definition of sighs, which unifies observations and conclusions in this and previous *in* vivo^6,17,19,20^ and *in vitro*^21,22^ studies that used various, often incongruent definitions of sighs.

Here, we investigated two novel properties of sigh generation: sigh phase and postsigh refractory period. By using the same approach that we employed for differentiating the effects of optogenetic stimulation on various preBötC subpopulations on the eupneic cycle^8,20,23^, we found that sighs evoked by optogenetic stimulation of various subpopulations reset the sigh cycle and delayed the onset of the next endogenous sigh. Also, similar to a postinpiratory refractory period in the respiratory cycle, during which ectopic breaths cannot be evoked immediately following termination of an inspiratory effort^8^, we similarly observed a postsigh refractory period following an endogenous sigh, during which ectopic sighs could not be evoked. We suggest that these subpopulations have intrinsic “sigh-generating” properties, e.g., unique signaling pathways or connectivity that affects excitability of these subpopulations. Although vagotomy transiently abolishes eupneic sighs^24^, they can be evoked in absence of sensory feedback *in vivo* after vagotomy^25^ or *in vitro*^21,22^, suggesting that sensory inputs by themselves are not necessary, but likely modulate activity of these subpopulations to affect frequency of sighs. The mechanisms that underlie the slow sigh rhythm (∼minutes) remains to be discovered, and might involve intracellular Ca^2+^ oscillations^26^.

In mice, sigh-like movements appear *in utero* between E17 and E18^27^, when at least two types of inspiratory-modulated neurons can be differentiated in rhythmic slices: neurons that fire in sync with regular “eupneic” XII bursts, and those that fire only during sigh-like bursts^27^. This could result from increased activity in a single population with a distribution of activation thresholds or from recruitment of a distinct high threshold population. *In vitro*, the majority of NMBR preBötC neurons recorded were active in both eupneic and sigh bursts, a few GRPR preBötC neurons recorded were active in eupnea, but none were active exclusively during sighs, q.v.,^27^. Rhythmic preBötC NMBR neurons in neonatal mice were active during both eupneic and sigh bursts (Fig. 4b), but not during burstlets or the pre-I period. These results suggest that NMBR preBötC neurons are Type II neurons that participate in generation of the burst patterns of sighs and eupneic breaths, but are not rhythmogenic.

Sighs can be induced by activation of various subpopulations within two structures: NMB or GRP in pF and NMBR, GRPR or SST in preBötC. We previously reported a microcircuit capable of generating sighs involving pF neurons producing bombesin-related peptides that project to preBötC neurons expressing cognate NMBRs and GRPRs^4^, and that activation of these preBötC neurons via their peptide receptors was sufficient to generate sighs^4^. Here, in significantly extending this work, we show that sighs can be evoked by direct photoexcitation of GRP, NMBR or GRPR neurons, bypassing activation of these receptors. Photostimulation of preBötC GRPR or NMBR neurons can elicit ectopic sighs (Fig. 3a and 3c), even in the presence of GRPR and NMBR antagonists at sufficient concentration to block the effect of exogenous application of peptides that increase sigh rate (Fig. 6d and Extended data Fig. 6). Thus, sighs can be induced by increasing the excitability of these subpopulations, without obligatory participation of pathways activated by bombesin-related peptides. In other words, while it can provide sufficient external input to trigger sighs, activation of preBötC NMBRs and/or GRPRs is not necessary for sigh production. This suggests that sighs are not the unique product of a preBötC bombesin-peptide signaling pathway.

We studied photoactivation of pF NMB- and GRP-expressing neurons in transgenic mice. Could our protocol have also activated GRP or NMB containing presynaptic terminals originating from brainstem neurons outside pF, such as found in the parabrachial nuclei and NTS^4^? This is unlikely in the case of NMB, since NMB-expressing neurons are limited to pF in the medulla^4^.

Also, unlikely the case of GRP, as there is no evidence that pF neurons express GRPRs^28^. However, our protocol would result in the expression of ChR2 in neurons that expressed NMB-or GRP-expressing neurons at any point since conception, even transiently. Thus, we cannot rule out that; in our adult mice there were neurons in pF that once expressed these peptides but no longer did so, and that these vestigial NMB or GRP neurons could have been perturbed in our protocol. Such determination requires additional studies outside the scope of the work reported here.

preBötC neurons expressing SST, mostly distinct from NMBR and GRPR populations (Fig. 2), receive input from rhythmogenic preBötC neurons, many have an inspiratory firing pattern^14^, that appear to serve a premotor role to shape inspiratory motor output^8^. Photostimulation of preBötC SST neurons increases peak inspiratory amplitude in both anesthetized and awake mice^8,23^. Here, DREADD-mediated excitation of SST neurons significantly increased sigh rate, with the effect much larger than DREADD-mediated excitation of NMBR neurons (16- vs 3-fold increase). Additionally, unlike the significant 1-5s latency to induce an ectopic sigh when photostimulating GRPR or NMBR preBötC neurons (Fig. 3e), photostimulating preBötC SST neurons during inspiration induced a second augmented inspiration immediately following the eupneic burst to convert it into a sigh (Fig. 7b). Importantly, effects of such photostimulation were unaffected by blockade of both NMBR and GRPR receptors (Fig.7d), and sighs evoked by microinjection of peptides NMB and GRP into preBötC were completely eliminated by inhibition of SST neurons that expressed an inhibitory DREADD PSAM4 (Fig. 7g and 7h). We conclude that activation of preBötC SST neurons is also sufficient to induce sighs and further suggest they can induce sighs, downstream from NMBR and GRPR preBötC neurons, and may be necessary to do so (Fig. 8).

**Fig. 8.**
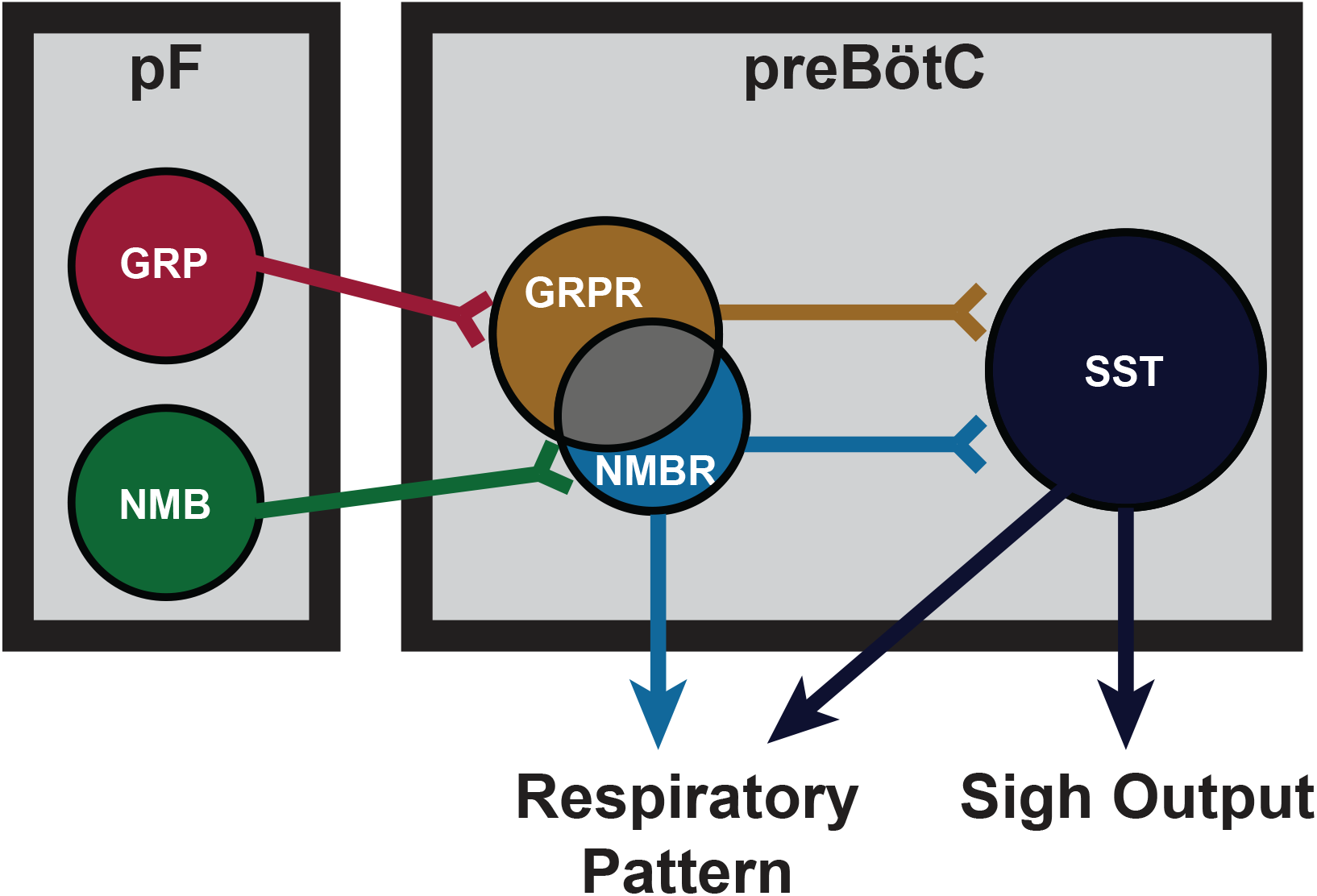
Proposed model of sigh generating pF-preBötC microcircuit. NMB and GRP neurons in pF project to preBötC neurons expressing cognate receptors NMBR and GRPR. NMBR and GRPR preBötC neurons mediate sigh output via connections to downstream SST preBötC neurons, but also directly contribute to the pattern of eupneic breathing.

Our initial delineation of a microcircuit of four subpopulations, i.e., pF NMB and GRP neurons and preBötC NMBR and GRPR neurons, that affected sighs led us to hypothesize its role in sigh generation, but we were agnostic as to its role in the production of eupneic breaths^4^. Our data here, consistent with a previous report of effects from intravenous administration of bombesin^29^, suggests that this microcircuit is tightly integrated within the eupneic CPG with NMBR, GRPR and SST populations having distinct though overlapping roles in modulation of the eupneic breathing pattern as well as in the generation of sighs. The majority of preBötC neurons expressing NMBR, GRPR (or both) receptors are glutamatergic and excitatory (Fig. 2). *In vitro*, no NMBR preBötC neurons were active during burstlets or the pre-I period, suggesting that NMBR preBötC neurons are Type II neurons that participate in generation of breathing pattern but not rhythm generation (similar to the properties of preBötC SST neurons^14^). *In vivo* we observed distinctly different effects from activation of preBötC GRPR and NMBR neurons: photoactivation of GRPR neurons increased breathing frequency, whereas photo- or chemo- activation of NMBR neurons decreased breathing frequency and increased amplitude (Fig. 6a and 6c); otherwise, their effects on sighing appeared similar (Fig. 3). Notably, some neurons express both NMBR and GRPR receptors (Fig. 2). When the overlapping subpopulation of preBötC neurons expressing both GRPRs and NMBRs was excluded, we could activate the two discrete pathways independently. Photostimulating GRPR-only neurons had no effect on inspiratory burst amplitude, whereas photostimulating NMBR-only neurons increased amplitude (Fig. 6b). Additionally, DREADD-mediated excitation of the SST population increased both the sigh rate and the inspiratory burst amplitude (Fig. 7f). Microinjection of peptides NMB and GRP into preBötC increased sigh rate and inspiratory burst amplitude; subsequent ultrapotent chemogenetic inhibition of preBötC SST^+^ neurons selectively abolished sighs, but not an increase in inspiratory amplitude (Fig. 7g and 7h), suggesting that modulation of breathing pattern by NMBR and GRPR neurons is independent of SST neurons (Fig. 8).

Sighing is a relatively straightforward behavior that appears to result from a delineable microcircuit that could only be discovered because of the close relationship between the essential, readily localizable, compact circuitry with a definitive precisely measurable functional physiologically relevant output. How this unexpected complexity for sigh generation reflects on strategies for unraveling ever more complex and difficult to quantify behaviors remains to be determined.

## Methods

Experimental procedures were carried out in accordance with the United States Public Health Service and Institute for Laboratory Animal Research Guide for the Care and Use of Laboratory Animals. All animals were handled according to institutional protocols at the University of California, Los Angeles (#1994-159-83P0) and approved by University of California Animal Research Committee (Animal Welfare Assurance #A3196-01). Every effort was made to minimize pain and discomfort, as well as the number of animals.

### Animals

Grp^Cre^ was obtained from Jackson Labs (Strain #033174). Grpr^flp^, Nmb^Cre^ and Nmbr^Cre^ animals were designed and generated by Cyagen Biosciences Inc (see below). Transgenic ChR2 mice were generated by crossing Cre mice with floxed-ChR2-tdTomato mice (Jackson Labs Strain #012567). These crosses generated mice expressing ChR2 in GRP neurons (Grp-ChR2), NMB neurons (Nmb-ChR2) or NMBR neurons (Nmbr-ChR2). To express ChR2 selectively in preBötC neurons, AAV injections were performed on adult Sst^Cre^ mice (Jackson Labs Strain #013044) or Grpr^flp^ mice. All *in vivo* experiments were performed on adult male or female mice (10-14 weeks old, 24-32g). Unless otherwise specified, mice were anesthetized with ketamine/xylazine. For *in vitro* experiments, transgenic Grpr-EGFP, Nmbr-tdTomato, and Nmbr-GCamp6s were used.

These mice were generated by crosses between Nmbr^Cre^ or Grpr^Flp^ mice and one of the following lines: Ai14 (Jackson Labs Strain #007914), RCE:FRT (MMRRC Strain #032038-JAX), GCaMp6f (Jackson Labs Strain #029626).

To generate Nmb^Cre^ mice the open reading frame of Cre recombinase together with SV40 polyA signal was cloned downstream of the mouse Nmb promoter, such that the expression of Cre mimicked the endogenous mouse Nmb gene. The PiggyBac ITRs was inserted into the BAC backbone flanking the genomic insert, to facilitate transposes mediated BAC integration. The modified BAC was co-injected with transposes into single cell stage fertilized eggs from C57BL/6 mice. Resulting pups were then genotyped by PCR for the presence of the modified BAC and the Cre cassette. To generate Grpr^Flp^ mice, the Grpr gene was first located on mouse chromosome X. Three exons have been identified, with the ATG start codon in exon 1 and TAG stop codon in exon 3; the TAG stop codon was then replaced with the Flp cassette. To engineer the targeting vector, homology arms were generated by PCR using BAC clone RP23-402H23 and RP23-50M18 from the C57BL/6J library as template. In the targeting vector, the Neo cassette was flanked by FRT sites. DTA was used for negative selection. The constitutive knock-in allele was obtained after FRT-mediated recombination. C57BL/6 ES cells were then used for gene targeting to introduce knock-in alleles into host embryos followed by transfer into surrogate mothers. Resulting pups were then genotyped by PCR for the presence of the Flp cassette. To generate Nmbr^Cre^ mice, the Nmbr gene was first located on mouse chromosome 10. Three exons have been identified, with the ATG start codon in exon 1 and TGA stop codon in exon 3; the TGA stop codon was then replaced with the Cre cassette. To engineer the targeting vector, homology arms was generated by PCR using BAC clone RP24-232I16 and RP24-124K10 from the C57BL/6J library as template. In the targeting vector, the Neo cassette was flanked by LoxP sites. DTA was used for negative selection. The constitutive knock-in allele was obtained after Cre-mediated recombination. C57BL/6 ES cells were then used for gene targeting to introduce knock-in alleles into host embryos followed by transfer into surrogate mothers. Resulting pups were then genotyped by PCR for the presence of the Cre cassette.

### Viral vector design

pAAV-hSyn Con/Foff hChR2(H134R)-EYFP (Addgene plasmid # 55646 ; http://n2t.net/addgene:55646 ; RRID:Addgene_55646) and pAAV-hSyn Coff/Fon hChR2(H134R)-EYFP (Addgene plasmid # 55648 ; http://n2t.net/addgene:55648 ; RRID:Addgene_55648) were gifts from Karl Deisseroth^15^. pAAV-hSyn-DIO-hM3D(Gq)-mCherry was a gift from Bryan Roth (Addgene plasmid # 44361; http://n2t.net/addgene:44361; RRID:Addgene_44361). pAAV-Syn-flex-PSAM4-GlyR-IRES-EGFP was a gift from Scott Sternson^16^ (Addgene plasmid # 119741; http://n2t.net/addgene:119741; RRID:Addgene_119741). All viruses were stored in aliquots at -80°C until use.

### Viral injections

Mice were anesthetized with isoflurane (4% for induction and 2% for maintenance) and placed in a stereotaxic apparatus (David Kopf Instruments) with Bregma and Lambda skull landmarks level. Two holes were drilled in the skull 6.80 mm caudal to Bregma and 1.2 mm lateral to the midline. Virus was delivered 4.65 mm from the dorsal surface of the brain into the preBötC through a glass pipette using either a pressure ejection system (Picospritzer II; Parker Hannafin) or Micro4 microinjection system (World Precision Instruments). For ChR2 experiments, we used 100-200 nl of virus solution (1-6 x 10^12^ vg/ml) per side, and for DREADD experiments, 30-50 nl of virus solution (4.6 x 10^12^ vg/ml) per side. The pipettes were left in place for 5 min after injection to minimize backflow. The wound was closed with 5-0 gauge non-absorbable sutures. Mice were returned to their home cage and given 2-3 weeks to recover to allow for sufficient levels of protein expression. All virus injection sites were subsequently confirmed by immunostaining and only mice with sufficient and localized transfection in preBötC were used for all analysis (see Extended Data Fig. 2 for representative confocal images).

### Surgical procedures for ventral approach

Adult mice were anesthetized via intraperitoneal injection of ketamine and xylazine (100 and 10 mg/kg, respectively). Isoflurane (1-2% volume in air) was administered throughout an experiment. The level of anesthesia was assessed by the suppression of the withdrawal reflex. A tracheostomy tube was placed in the trachea through the larynx, and respiratory flow was detected with a flow head connected to a transducer to measure airflow (Mouser Electronics). The mice were placed in a supine position in a stereotaxic instrument (David Kopf Instruments). The larynx was denervated, separated from the pharynx, and moved aside. The basal aspect of the occipital bone was removed to expose the ventral aspect of the medulla. The canal of the hypoglossal nerve (XII) served as a suitable landmark. The preBötC were 0.15 mm caudal to the hypoglossal canal, 1.2 mm lateral to the midline, and 0.24 mm dorsal to the ventral medullary surface.

### Photostimulation

A 473 nm laser (OptoDuet Laser; IkeCool) was targeted bilaterally to the preBötC or pF with a branching fiber patch cord (200 µm diameter; Doric Lenses) brought to the exposed ventral medullary surface. Laser power was set at 5 mW. Short Pulse Photostimulation (SPP; 200 ms) and Long Pulse Photostimulation (LPP; 5 s) waveforms were delivered under the command of a pulse generator (Pulsemaster A300 Generator; WPI) connected to the laser power supply. In instances where breathing frequency was decreased, e.g. after BIM23042 and RC3095 microinjections, SPP and LPP of longer durations (500 ms and 8 s, respectively) were used to confirm consistency of results. Stimulating 500 µm rostral to preBötC or pF in mice expressing ChR2 that served as a control did not produce significant output effects (Extended data Fig. 7b). No output effects and no sigh were produced by preBötC or pF photostimulation in Grp-RFP, Grpr-EGFP, Nmbr-EGFP, Nmb-RFP reporter mice (Extended data Fig. 7a).

### Pharmacological injection experiments

The NMBR antagonist BIM23042 (Tocris Bioscience) and the GRPR antagonist RC3095 (Sigma-Aldrich) were injected together (300 µM each, 50-60 nl/side) to block NMBR and GRPR activation. Injections were made using micropipettes (∼40 µm tip), placed bilaterally into the preBötC. Injections targeted to the center of the preBötC were placed 0.15 mm caudal to the hypoglossal canal, 1.2 mm lateral to the midline, and 0.24 mm dorsal to the ventral medullary surface^8^. Small corrections were made to avoid puncturing of blood vessels on the surface of the medulla. All injections were made using a series of pressure pulses (Picospritzer, Parker- Hannifin; or Micro4, World Precision Instruments). For activation of hM3Dq receptors, 50µL of CNO solution was applied to the ventral surface of the brainstem. CNO (Tocris Biosciences) was dissolved in saline at concentration of 1 mg/ml with 0.5% dimethylsulfoxide. Solution was prepared fresh daily. For activation of PSAM4-GlyR receptors, 30µL of uPSEM817 tartrate solution (10mM) was applied to the ventral surface of the brainstem. In chemogenetic experiments CNO or uPSEM817 were applied to the ventral surface of the brainstem instead of microinjection in order to avoid preBötC damage from multiple microinjections; in control experiments application of the same dose of CNO or uPSEM817 does not produce responses in any breathing parameters.

### *In situ* hybridization and immunostaining

Nmbr^Cre^, SST^Cre^ or wild-type mice (10-14 weeks) were euthanized with isoflurane overdose, their brainstems were rapidly removed and flash frozen in dry ice. Fresh frozen brainstems were sectioned sagittally on a cryostat (CryoStar NX70, Thermo Scientific), mounted on SuperFrost Plus slides (Fisher Scientific) and stored at -80 °C for at least 1 day. They were then processed according to manufacturer’s protocol (RNAscope version 1, Advanced Cell Diagnostics). Briefly, tissue samples were postfixed in 10% neutral buffered paraformaldehyde, washed, and dehydrated in sequential concentrations of ethanol (50, 70, and 100%). Samples were treated with protease IV and incubated for 2 hours at 40 °C in HybEZ^TM^ Hybridization Oven (Advanced Cell Diagnostics) in the presence of target probes. We used combinations of following probes for hybridization: *Mm-ChAT*, *Mm-Grpr, Mm-Nmbr, Mm-Sst*, *Mm-Slc17a6 (VGluT2), Mm-Aldh1l1, Cre* and *Flp*. A probe for *Mm-ChAT* was used in all probe combinations to mark the location of nucleus ambiguus, which is necessary for determining the location of preBötC. After a 4-step amplification process, samples were counterstained with DAPI and coverslipped with ProLong Gold (Invitrogen) used as a mounting agent. Images were acquired on a confocal laser scanning microscope (LSM710 META, Zeiss). High-resolution z-stack confocal images were taken at 1 µm intervals and then merged using ImageJ software.

Immunohistochemistry was performed according to the following protocol. Free-floating sections were rinsed in PBS and incubated with 10% normal donkey antiserum (NDS) and 0.2% Triton X-100 in PBS for 60 min to reduce nonspecific staining and increase antibody penetration. Sections were incubated overnight with primary antibodies diluted in PBS containing 1% NDS and 0.2% Triton X-100. The following day, sections were washed in PBS, incubated with the specific secondary antibodies conjugated to the fluorescent probes diluted in PBS for 2 h. Sections were further washed in PBS, mounted, and coverslipped with Fluorsave mounting medium (Millipore). The primary antibodies used for this study were as follows: rabbit polyclonal anti-somatostatin-14 (1:500; Peninsula Laboratories), mouse monoclonal anti-NeuN (1:500; Millipore; MAB377), goat anti-ChaT (1:500, Chemicon) and chicken polyclonal anti-GFP (1:500; Aves Labs). DyLight488 donkey anti-chicken, Rhodamine Red-X donkey anti-rabbit, and Cy5 donkey anti-mouse conjugated secondary antibodies (1:250; Jackson ImmunoResearch) were used to detect primary antibodies. Slides were observed under an AxioCam2 Zeiss fluorescent microscope connected with AxioVision acquisition software or under a LSM510 Zeiss confocal microscope with Zen software (Carl Zeiss). Images were acquired, exported in TIFF files, and arranged to prepare final figures in Zen software (Carl Zeiss) and Adobe Photoshop (Adobe).

### *In vitro* electrophysiology

Neonatal mice (P0-5) of either sex were anesthetized with isoflurane. The brainstem was isolated from the pons to the rostral cervical spinal cord under cold ACSF containing (in mM): 124 NaCl, 3 KCl, 1.5 CaCl_2_, 1 MgSO_4_, 25 NaHCO_3_, 0.5 NaH_2_PO_4_, and 30 D-glucose; a single transverse slice (600μm) was cut on a vibratome (Leica VT1200S).

Slices were placed in a recording chamber with the rostral face of preBötC facing up and continuously superfused with warmed ACSF (at 28-30°C) equilibrated with 95% O_2_ and 5% CO_2_, and extracellular K^+^ was increased to 9 mM to elevate preBötC excitability. Inspiratory- related motor output was recorded from the hypoglossal nerve (XII) using a suction electrode and a differential AC amplifier (AM systems), filtered at 2-4 kHz, integrated, and digitized at 10 kHz. Whole-cell recordings were made using a MultiClamp 700A (Molecular Devices), filtered at 2-4 kHz, and digitized at 10 kHz. Borosilicate glass recording electrodes (G150-4, Warner Instruments) had tip resistances of 4-8 MΩ and were filled with intracellular solution containing (in mM): 140 potassium gluconate, 10 HEPES, 5 NaCl, 1.1 EGTA, 2 Mg-ATP, and 0.1 CaCl_2_ (∼305 mOsm; pH 7.3). In two-photon imaging experiments, field potentials were recorded from the contralateral preBötC, using an extracellular electrode and a differential AC amplifier (AM systems), filtered at 2-4 kHz, integrated, and digitized at 10 kHz. Digitized data were analyzed off-line using custom procedures written for IgorPro (Wavemetrics). Agonists were bath applied at the specified concentrations while monitoring field potentials and XII output.

### Two-photon calcium imaging

Intracellular Ca^2+^ was imaged using a multi-photon resonant scanner (3i) and a microscope equipped with a water immersion 20x, 1.0 objective. Illumination was provided by a laser with a power output of 1050 mW at 970 nm (Coherent Chameleon UltraI). Identified preBötC neurons were scanned at 10-39 Hz and fluorescence data were collected using 3i software and analyzed using SlideBook, Image J and IgorPro. Regions of interest (ROIs) were manually detected and Ca^2+^ transients were plotted as F/F_0_, where F_0_ is the average fluorescence intensity of all pixels within a given ROI averaged over the entire time series.

### Statistical analysis

*In vitro* data were analyzed using IgorPro (Wavemetrics) and GraphPad Prism (GraphPad Software) was used for statistical analysis. Paired T-tests or Kruskal-Wallis one-way ANOVA, adjusted for multiple comparisons, was used to calculate significance.

For *in vivo* experiments, data were recorded on a computer using LabChart 7 Pro (AD Instruments) and analyzed using LabChart 7 Pro (ADInstruments), Excel, and Igor Pro (Wavemetrics, Inc.) software. The flow signal was high-pass filtered (> 0.1 Hz) to eliminate DC shifts and slow drifts, and was digitally integrated with a time constant of 0.05 s to calculate breathing frequency, period, V_T_, *T_I_* and *T_E_* during eupnea or sigh (Extended data Fig. 1d).

Breathing frequency is measured by the time interval between peaks. *T_I_* was calculated by the time interval from start to the maximum of the peak, *T_E_* was calculated by the time interval from the previous inspiratory peak to the onset of the next inspiratory burst and V_T_ was calculated from the inspiratory peak. Due to high reproducibility of effects of photostimulation, V_T_, *T_I_* and *T_E_* during ectopic sighs were calculated as an average of 3-5 sighs, paired t-tests were used to determine statistical significance of changes before and after pharmacological injections or photostimulation. For statistical comparisons of more than two groups, repeated-measures (RM) ANOVAs were performed. For one-way RM ANOVAs, post hoc significance for pairwise- comparisons was analyzed using Holm-Sidak method. Significance was set at p < 0.05. Data are shown as mean ± SE.

## Data availability

The data that support the findings of this study are available from the corresponding author, JLF, upon reasonable request.

## Author Contributions

J.L.F., Y.C., E.B., C.T.P., and D.N.C. conceived experiments, interpreted data, and wrote the manuscript. Y.C. performed experiments showing the effects on sighing and other aspects of breathing pattern of optogenetic activation of the pF GRP or NMB neurons and preBötC GRPR, NMBR or SST neurons *in vivo*. E.B. performed experiments showing the effects on sighing and other aspects of breathing pattern of chemogenetic excitation and inhibition of preBötC NMBR or SST neurons *in vivo*. C.T.P. performed two-photon calcium imaging experiments showing the activity profiles of preBötC NMBR neurons *in vitro*. D.N.C. performed whole-cell recordings of preBötC NMBR and GRPR neurons *in vitro.* E.B. performed the RNAscope experiments. Y.C., E.B., C.T.P., and D.N.C. analyzed data from their respective experiments.

## Acknowledgements

This work was funded by NIH Grants HL135779, NS72211, NSFC Grant 37101012 and Disciplinary Construction Innovation Team Foundation of Chengdu Medical College: CMC-XK-2102.

---

**Extended data Fig. 1.**
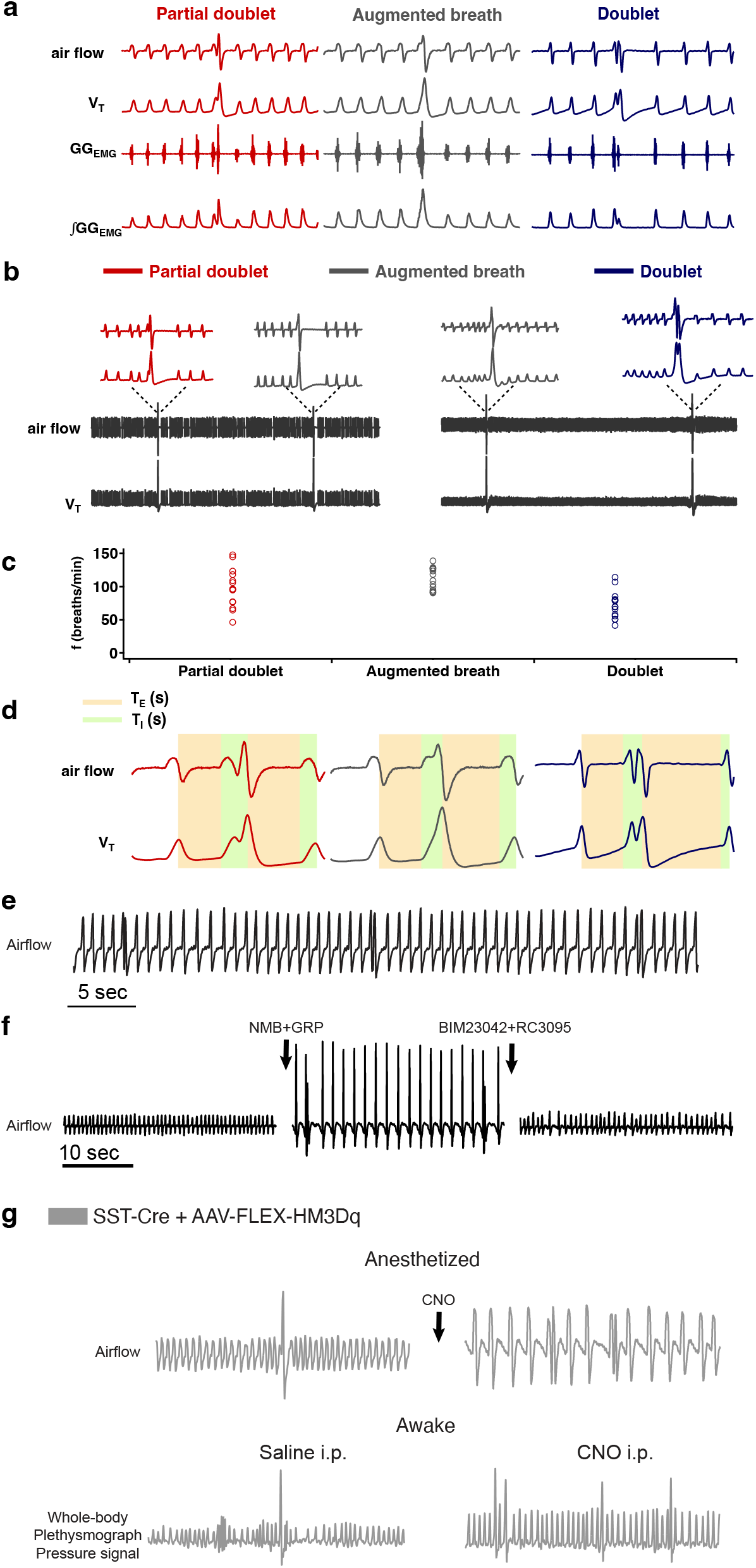
a, Three types of sighs observed in adult mice *in vivo*. A sigh is a spontaneous inspiratory effort that results in significantly increased inspiratory tidal volume, typically two to five times compared to normal breaths. Under anesthesia in mice, sighs *in vivo* can take multiple shapes. The most common shape is partial doublet, a biphasic double-sized breath with an initial phase that is identical to a normal breath (eupnea) and a later high-amplitude inspiration, coincident with a biphasic genioglossus_EMG_ (GG_EMG_) event (left). An *in vivo* sigh in mice can also present as one large breath, a monophasic augmented inspiration that has two to five times the volume of a normal breath, coincident with a monophasic GG_EMG_ event (middle); or a double-peaked breath (doublets) with the first breath immediately followed by a similar amplitude second breath (right). **b**, Raw and expanded traces show three types of shape observed in one mouse. **c**, Sigh shapes are related to basal breathing frequency (45 spontaneous sighs from 3 mice). Different sigh shapes represent a continuum from “augmented breath” on one side of the spectrum to “doublet” on the other. Exact shape appears to be related to basal breathing frequency, with augmented breaths and occasional partial doublets exhibited during conditions with high basal respiratory rate/low amplitude; while doublets and partial doublets occur mostly with low basal respiratory rate/high amplitude. The latter is similar to conditions of vagotomy and *in vitro*, where breathing *f* is substantially reduced, and during which sighs often appeared as doublets. **d**, Measurement of respiratory parameters of eupneic breath and three types of sighs. **e,** Sighs exhibiting a doublet shape occur periodically at a frequency significantly lower than normal eupneic breaths after NMB and GRP microinjection into preBötC in ketamine/xylazine anesthetized mice. **f,** Doublets are not artefacts resulting from any damage to preBötC, since doublet-shaped sighs induced by microinjection of NMB and GRP were blocked by subsequent preBötC microinjection of NMBR and GRPR antagonists BIM23042 and RC3095. **g,** Chemogenetic excitation of SST preBötC neurons evokes frequent doublet-shaped sighs in ketamine/xylazine mice, while in awake mice it induces frequent sighs of augmented breath shape, suggesting that doublets are indeed sighs.

**Extended data Fig. 2.**
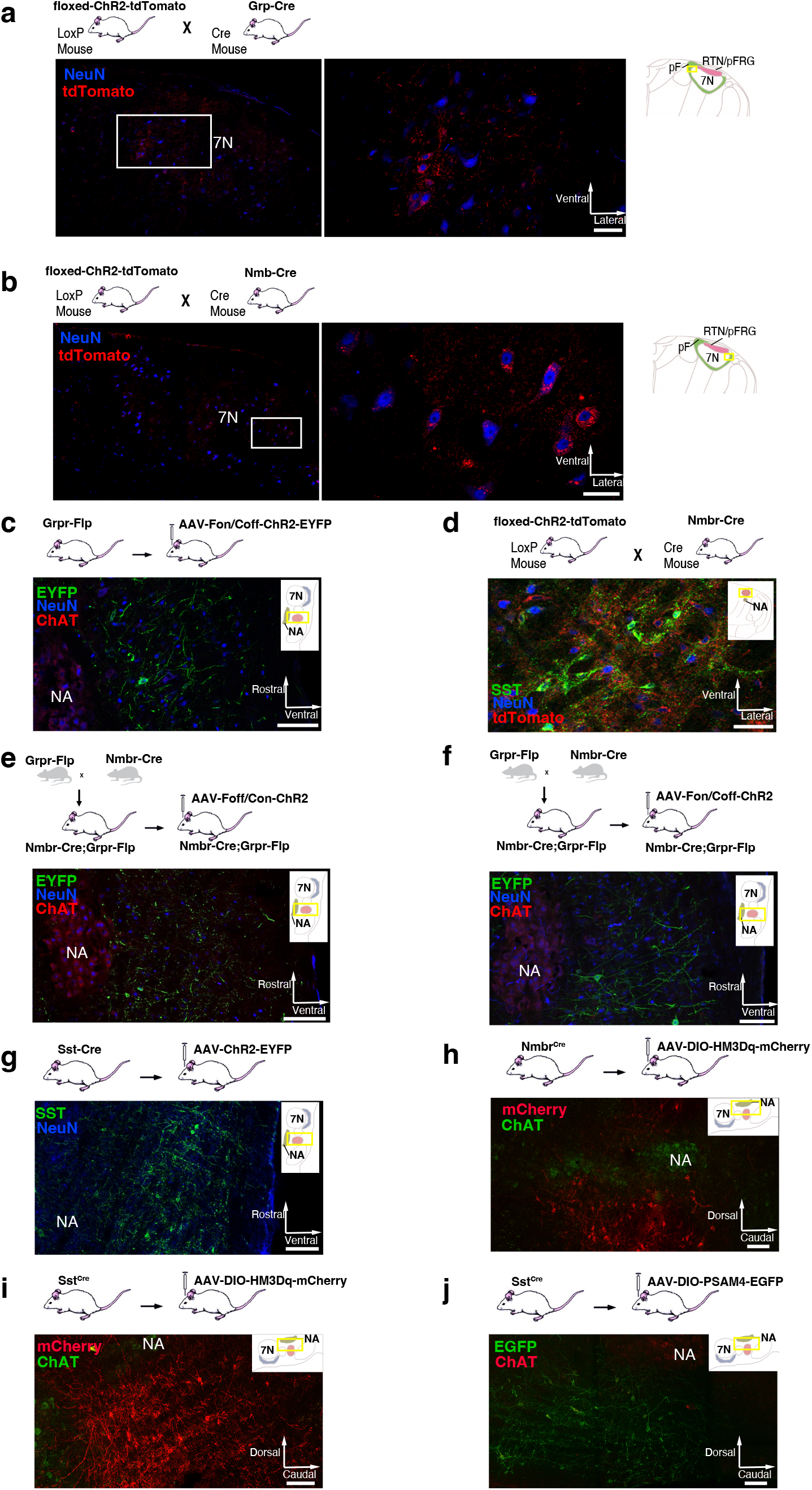
a, tdTomato (red) in the pF neurons (blue) of Grp-ChR2 mice. Left: Coronal brainstem section at the level of pF. Right: High-magnification micrographs of square segment in left panel show ChR2 expression (tdTomato, red) in pF GRP neurons. Section also labeled for NeuN (blue) immunoreactivity. Scale bar, 50 µm. **b,** tdTomato (red) in the pF neurons (blue) of Nmb-ChR2 mice. Left: Coronal brainstem section at the level of pF. Right: High-magnification micrographs of square segment in left panel show ChR2 expression (tdTomato, red) in pF NMB neurons. Section also labeled for NeuN (blue) immunoreactivity. Scale bar, 50 µm. **c,** Representative confocal micrograph of sagittal medullary section (rectangle segment in inset panel) of Grpr^Flp^ mice at the level of preBötC shows Flp-dependent ChR2 expression (EYFP, green) targeted to preBötC GRPR neurons. ChR2-EYFP expression was detectable in preBötC neurons 4 weeks after viral injection, with some expression in neuron axons outside the preBötC including in the BötC rostral to the preBötC; in the intermediate reticular formation (IRt) ventral to the Amb. We found very few ChR2-EYFP expressed axons in neighboring ventral respiratory column (VRC) caudal to the preBötC, or at more distant brainstem sites, e.g., 7N. No evidence of transfected neuron soma was found outside the preBötC. Section also labeled for ChAT (red) and NeuN (blue) immunoreactivity. Scale bar, 100 µm. **d,** Representative confocal micrograph of coronal medullary section (rectangle segment in inset panel) of transgenic Nmbr-ChR2 mice at the level of preBötC shows ChR2 expression (tdTomato, red) in preBötC NMBR neurons. ChR2- tdTomato expression was detectable in the preBötC neurons, plus scattered neurons outside the preBötC including in the intermediate reticular formation (IRt) ventral to the Amb and the gigantocellular reticular nucleus (Gi) dorsomedial to the preBötC. Section also labeled for SST (green) and NeuN (blue) immunoreactivity. Scale bar, 50 µm. **e,** Representative confocal micrograph of sagittal medullary section (rectangle segment in inset panel) of Nmbr^Cre^;Grpr^Flp^ double mutant mice at the level of preBötC shows ChR2 expression (EYFP, green) targeted to preBötC NMBR-only neurons. ChR2-EYFP expression was detectable in preBötC neurons 4 weeks after viral injection, with some expression in neuron axons in the BötC. No evidence of transfected neuron soma was found outside the preBötC. Section also labeled for ChAT (red) and NeuN (blue) immunoreactivity. Scale bar, 100 µm. **f,** Representative confocal micrograph of sagittal medullary section (rectangle segment in inset panel) of Nmbr^Cre^;Grpr^Flp^ double mutant mice at the level of preBötC shows ChR2 expression (EYFP, green) targeted to preBötC GRPR-only neurons. ChR2-EYFP expression was detectable in preBötC neurons 4 weeks after viral injection, with very little expression in neuron axons in neighboring area outside the preBötC. No evidence of transfected neuron was found outside the preBötC. Section also labeled for ChAT (red) and NeuN (blue) immunoreactivity. Scale bar, 100 µm. **g,** Representative confocal micrograph of sagittal medullary section (rectangle segment in inset panel) of Sst^Cre^ mice at the level of preBötC shows Cre-dependent ChR2 expression (EYFP, green) targeted to preBötC SST neurons. ChR2-EYFP expression was detectable in preBötC neurons 4 weeks after viral injection, with some expression in neighboring neurons outside the preBötC including in the BötC rostral to the preBötC. The peak density of transfected neurons was caudal to the BötC and ventral to the Amb. We found no evidence of transfected neurons at more distant brainstem sites. Section also labeled for NeuN (blue) immunoreactivity. Scale bar, 100 µm. **h,** Representative confocal micrograph of sagittal medullary section (rectangle segment in inset panel) of Nmbr^Cre^ mice at the level of preBötC shows Cre-dependent HM3Dq expression (mCherry, red) targeted to preBötC NMBR neurons. HM3Dq-mCherry expression was detectable in preBötC neurons 4 weeks after viral injection, with the highest density of transfected somas in the preBötC. No evidence of transfected neuron was found outside preBötC. Section also labeled for ChAT (green) immunoreactivity. Scale bar, 100 µm. **i,** Representative confocal micrograph of sagittal medullary section (rectangle segment in inset panel) of SST^Cre^ mice at the level of preBötC shows Cre-dependent HM3Dq expression (mCherry, red) targeted to preBötC SST neurons. HM3Dq-mCherry expression was detectable in preBötC neurons 4 weeks after viral injection, with the highest density of transfected somas in the preBötC and minimal (<10% of transfected somas) found outside of preBötC. Section also labeled for ChAT (green) immunoreactivity. Scale bar, 100 µm. **j,** Representative confocal micrograph of sagittal medullary section (rectangle segment in inset panel) of SST^Cre^ mice at the level of preBötC shows Cre-dependent PSAM4 expression (EGFP, green) targeted to preBötC SST neurons. PSAM4-EGFP expression was detectable in preBötC neurons 4 weeks after viral injection, with the highest density of transfected somas in the preBötC and minimal (<10% of transfected somas) found outside of preBötC. Section also labeled for ChAT (red) immunoreactivity. Scale bar, 100 µm.

**Extended data Fig. 3.**
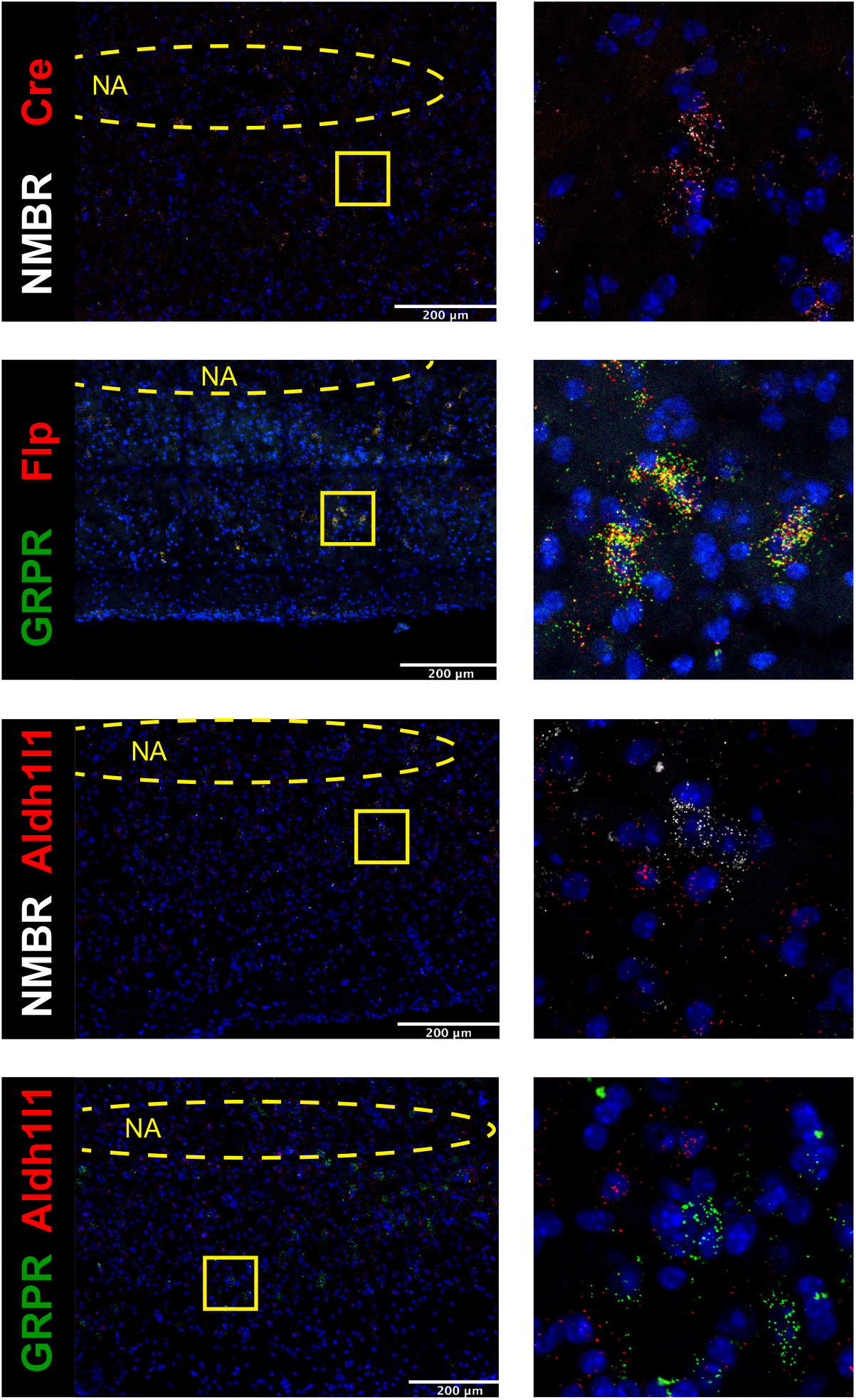
Colocalization between NMBR and Cre (A) and GRPR and Flp (B). All NMBR neurons expressed transcripts for Cre in brainstem tissue from Nmbr^Cre^ mice and all GRPR neurons expressed transcripts for Flp in brainstem tissue from Grpr^Flp^ mice. Colocalization between transcripts for NMBR, GRPR and astrocytic marker Aldh1l1 (C-D). Aldh1l1 was expressed throughout the brain tissue, with no clear boundaries demarking cells; therefore, we used NMBR and GRPR expression to determine cells boundaries and then assessed presence of Aldh1l1 within those boundaries. Vast majority of NMBR^+^ and GRPR^+^ neurons (97%) did not contain transcripts for Aldh1l1. NA: Nucleus ambiguous, location was determined using ChAT probe. Yellow squares indicate zoomed images (right, 100x100µm)

**Extended data Fig. 4.**
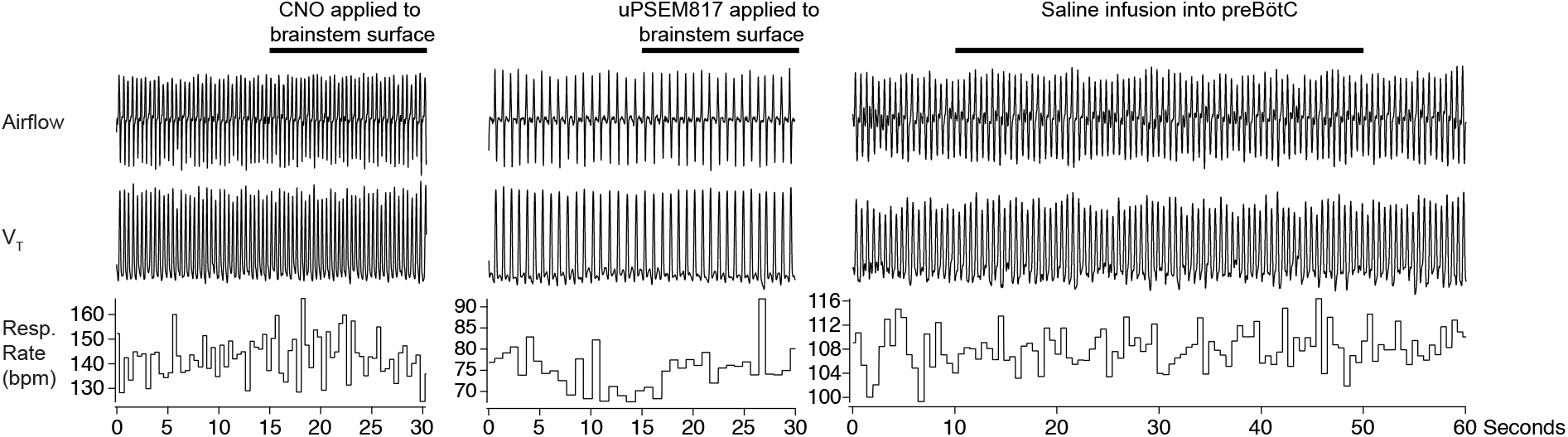
Control applications of CNO, uPSEM817 or Saline. We confirmed that application of CNO (left), uPSEM817 (middle) or saline infusion into preBötC did not evoke changes in breathing parameters and do not evoke sighing.

**Extended data Fig. 5.**
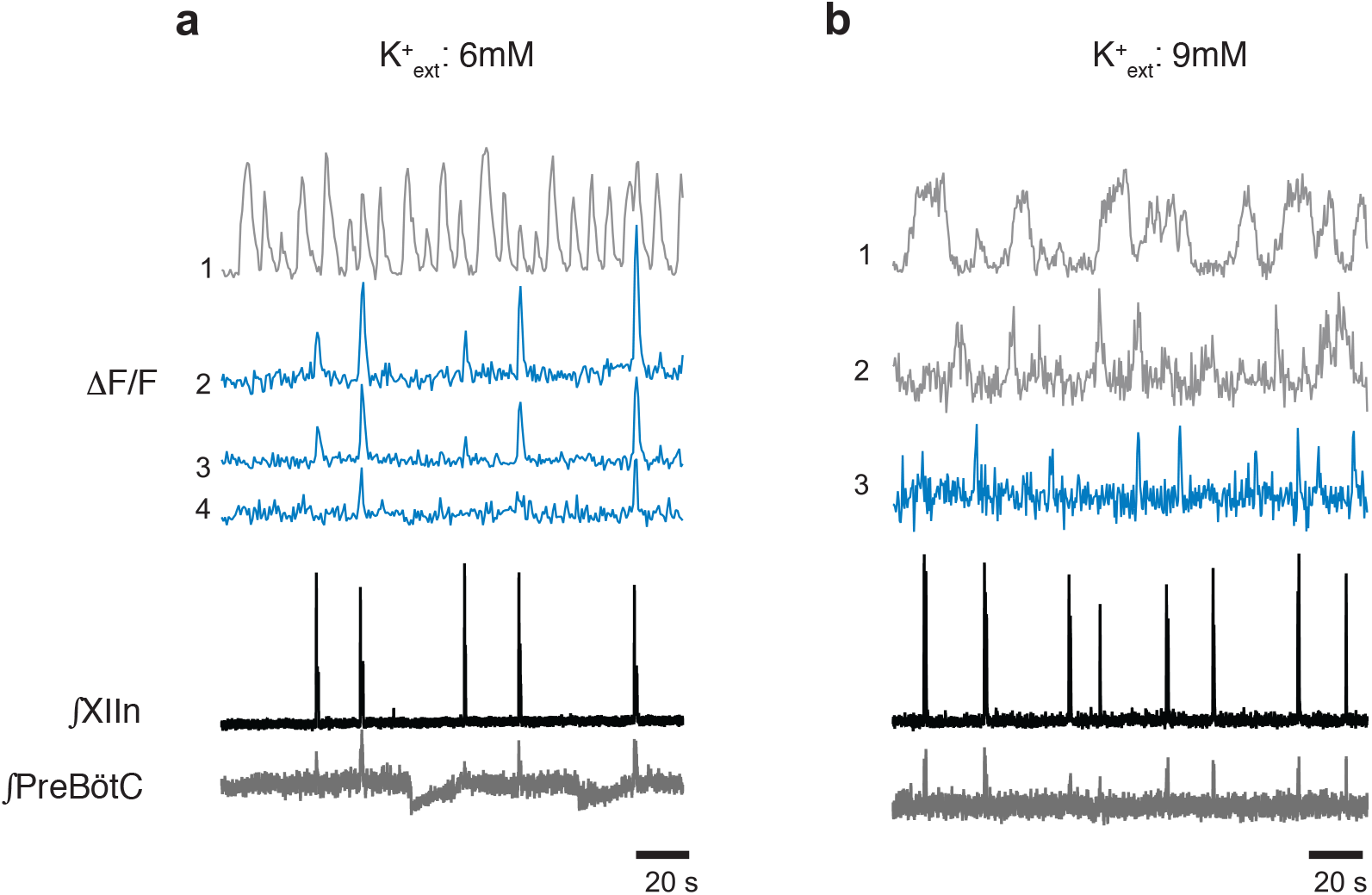
Examples of Ca^2+^ oscillations *in vitro* in phase with inspiratory rhythm (blue) and out of phase (gray) in a) 6 mM [K^+^]_ext_ (four neurons) and b) 9 mM [K^+^]_ext_ (three neurons).

**Extended data Fig. 6.**
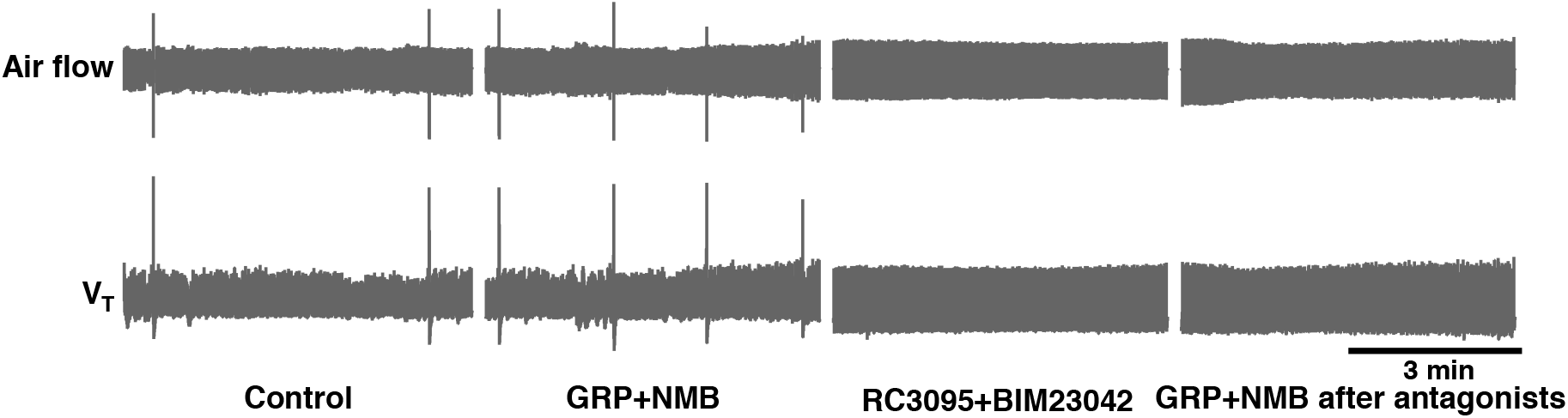
GRP and NMB receptors in preBötC effectively blocked by RC3095+BIM23042 microinjections. RC3095 and BIM23042 effectively blocked GRPRs and NMBRs, as respiratory responses induced by GRP and NMB microinjection into the preBötC, sufficient to increase sigh rate by 585 ± 209% of control, were fully antagonized by the same concentration of RC3095 and BIM23042 (n = 3). Air flow and V_T_ traces show the changes in sigh rate under different conditions (from left to right) in anesthetized mice (note sighs are evident as single events significantly larger than normal breaths): control; GRP+NMB: after bilateral injection of GRP+NMB into preBötC; RC3095+BIM23042: after bilateral injection of RC3095+BIM23042 (300 µM each, 50-60 nl/side) into preBötC; GRP+NMB after antagonists: the second bilateral injection of GRP+NMB, i.e., same dose of GRP+NMB that resulted in the higher sigh rate before RC3095+BIM23042 did not induce sighing after the antagonist injection.

**Extended data Fig. 7.**
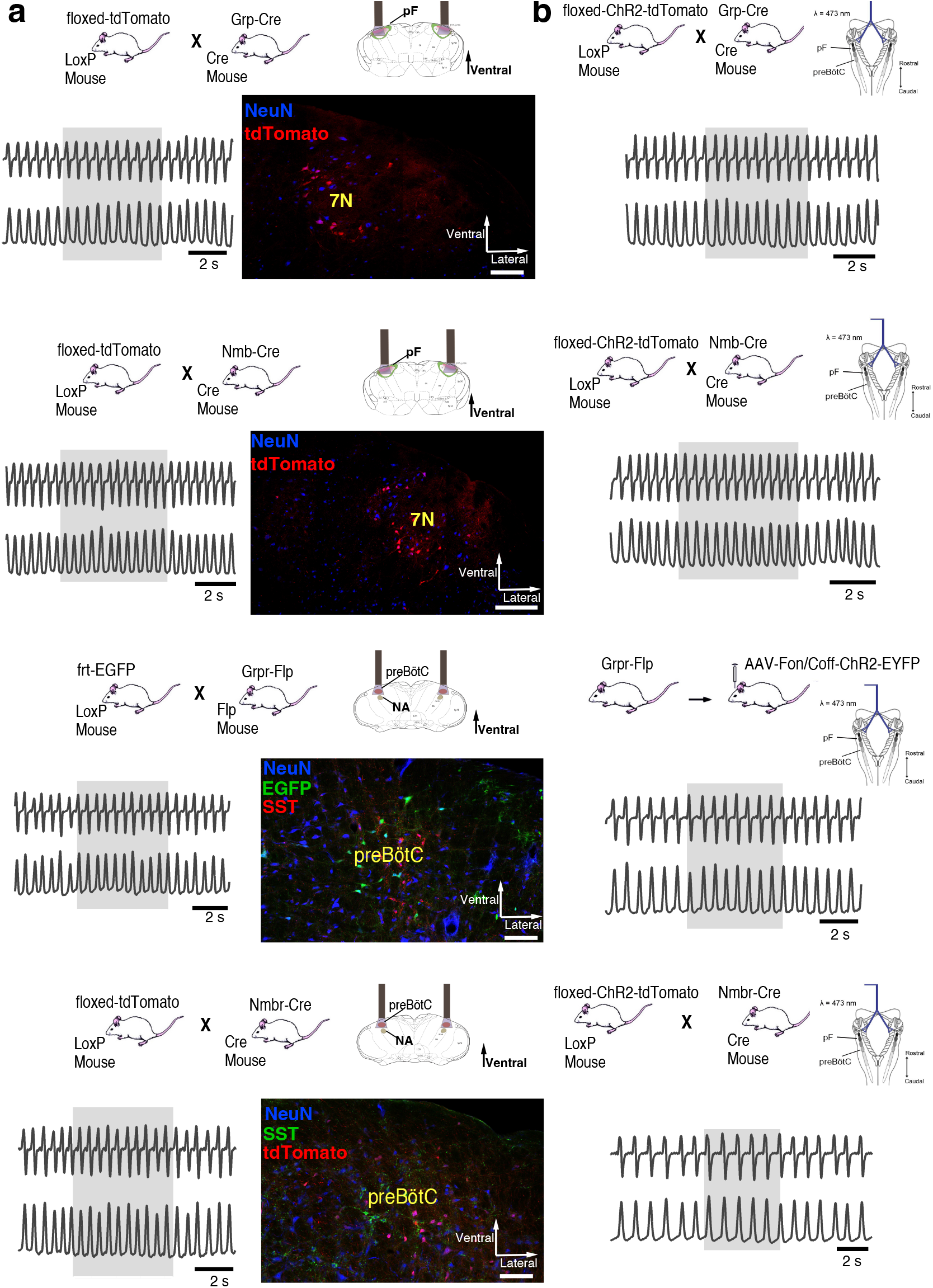
a, No output effects and no sigh were produced by pF or preBötC photostimulation in (from top to bottom) Grp-RFP, Nmb-RFP, Grpr-EGFP and Nmbr-RFP reporter mice. **b,** Stimulating 500 µm rostral to pF or preBötC in ChR2 expressed mice (from top to bottom: Grp-ChR2, Nmb-ChR2, Grpr-Flp and Nmbr-ChR2 mice) did not produce significant output effects.

